# BRAF^V600E^ induces reversible mitotic arrest in human melanocytes via microRNA-mediated suppression of AURKB

**DOI:** 10.1101/2020.05.21.109397

**Authors:** Andrew S. McNeal, Rachel L. Belote, Hanlin Zeng, Marcus Urquijo, Kendra Barker, Rodrigo Torres, Meghan Curtin, A. Hunter Shain, Robert H. I. Andtbacka, Sheri L. Holmen, David H. Lum, Timothy H. McCalmont, Matthew W. VanBrocklin, Douglas Grossman, Maria L. Wei, Ursula E. Lang, Robert L. Judson-Torres

## Abstract

Benign melanocytic nevi frequently emerge when an acquired *BRAF*^*V600E*^ mutation triggers unchecked proliferation and subsequent arrest in melanocytes. Recent observations have challenged the role of oncogene-induced senescence in melanocytic nevus formation, necessitating investigations into alternative mechanisms for the establishment and maintenance of proliferation arrest in nevi. We compared the transcriptomes of melanocytes from healthy human skin, nevi, and melanomas arisen from nevi and identified a set of microRNAs as highly expressed nevus-enriched transcripts. Two of these microRNAs – MIR211-5p and MIR328-3p – induced mitotic failure, genome duplication and proliferation arrest in human melanocytes through convergent targeting of AURKB. We demonstrate that *BRAF*^*V600E*^ induces a similar proliferation arrest in primary human melanocytes that is both reversible and conditional. Specifically, *BRAF*^*V600E*^ expression stimulates either arrest or proliferation depending on the differentiation state of the melanocyte. We report genome duplication in human melanocytic nevi, reciprocal expression of AURKB and microRNAs in nevi and melanomas, and rescue of arrested human nevus cells with AURKB expression. Together, our data describe an alternative molecular mechanism for melanocytic nevus formation that is congruent with both experimental and clinical observations.

## Introduction

Cutaneous melanoma is a potentially deadly skin cancer arising from the pigment-producing melanocytes of the human epidermis. An activating mutation in the BRAF proto-oncogene (BRAF^V600E^) drives over half of all cutaneous melanomas^1–3^. Yet, when a melanocyte acquires a BRAF^V600E^ mutation, the cell does not immediately transform to cancer. Instead, it usually undergoes clonal proliferation followed by stable arrest resulting in a benign skin lesion known as a melanocytic nevus or “mole”^3–6^. Despite the continued expression of BRAF^V600E^, the majority of melanocytic nevi remain innocuous for the lifespan of the individual, suggesting that nevus cells have robust intrinsic defenses against hyperproliferation. By inducing hyperproliferation and subsequent arrest, the BRAF^V600E^ mutation elicits divergent, biphasic phenotypes within a single cell – a poorly understood phenomenon. Characterization of the mechanisms and environmental factors that distinguish the two phenotypes could illuminate candidate strategies for chemoprevention of melanocytic nevus formation or interception of melanoma initiation.

The BRAF^V600E^ mutation is clonally present in ∼80% of human melanocytic nevi^3^ and melanocyte-specific expression of the oncogene is sufficient to induce nevus formation in mice^6–8^, suggesting that BRAF^V600E^ drives proliferation arrest in melanocytes. The prevailing theory to explain this seemingly paradoxical role for BRAF^V600E^ is oncogene-induced senescence (OIS)^9^. Cellular senescence is defined as the permanent cell-cycle arrest of previously replication-competent cells^10,11^. Senescence is associated with a variety of molecular hallmarks including elevated expression of p16^INK4A^ and other cyclin-dependent-kinase inhibitors, TP53, H2AX, lysosomal beta-galactosidase and senescence-associated secretory proteins (SASP) as well as a DNA content profile indicative of arrest in the G0/G1 phases of the cell cycle^11–16^. The concept of OIS as a barrier to tumorigenesis derived from observations of a durable proliferation arrest induced by overexpression of oncogenic RAS or RAF in immortalized human fibroblasts^12,17^.

More recent studies have questioned whether the term “senescence” aptly describes proliferation arrest of melanocytic nevi. Cell cycle re-entry that accompanies nevus recurrence^18^, eruption^19^, or transformation to primary melanoma suggests that the nevus arrest phenotype is reversible rather than permanent. Similarly, the expression of senescence markers do not readily distinguish human melanocytic nevi from primary or transformed melanocytes in humans (16^INK4A^, TP53, H2AX and beta-galactosidase) or mice (any known hallmark of senescence)^8,20,21^. Setting aside the permanence of OIS, melanocytic nevus arrest was largely thought to be driven by induction of p16^INK4A^ expression from the *CDKN2A* gene. Indeed, early studies demonstrated that BRAF^V600E^ induces p16^INK4A^ expression and that melanocytic nevi frequently express high levels of p16^INK4A 9^. Despite this, p16^INK4A^ expression is neither ubiquitous across melanocytic nevi, nor uniform within the cells of a single nevus^22,23^, and neither knock down of the 16^INK4A^ transcript nor ablation of the *CDKN2A* locus prevents the onset or rescues BRAF^V600E^-induced proliferation arrest^9,23–25^. BRAF^V600E^ also stimulates expression of p15^INK4B^ - the translational product of the *CDKN2B* gene^25^ which is situated adjacent to the *CDKN2A* locus - suggesting BRAF^V600E^ might orchestrate arrest via multiple cyclin dependent kinase inhibitors that induce arrest in the G1 phase of the cell cycle, prior to genome synthesis. While the loss of the *CDKN2A/B* locus is a formative event in melanoma progression, genetic evolution studies of melanoma progression suggest that selection against the *CDKN2A/B* locus occurs after BRAF^V600E^ melanocytes have already escaped nevus-associated arrest^23,26^. Collectively, these data complicate the straightforward model wherein BRAF^V600E^ induces the expression of cell cycle regulators leading to proliferation arrest, which is later circumvented by the genetic loss of those regulators.

Given these observations, we reasoned that the arrest response of human melanocytes to BRAF^V600E^ might be distinct from previous reports of BRAF^V600E^-induced OIS in other cell types. Specifically, we hypothesized that nevus-associated arrest is conditional and coordinated by reversible changes in gene expression. Here, we identify a transcriptional program that is consistently elevated in melanocytic nevi and includes microRNAs (miRNAs) previously shown to have diagnostic value^27,28^. In parallel, we observe that, unlike RAF-induced OIS in fibroblasts, BRAF^V600E^–induced arrest in human melanocytes is reversible and conditional. Merging these two observations, we discover that in melanocytes, regulation of miRNAs coupled to differentiation status permits toggling between BRAF^V600E^-driven hyperproliferative to arrested phenotypes.

## Results

### Melanocytic nevus-enriched miRNAs induce mitotic failure and proliferation arrest in primary human melanocytes

Since melanocytic nevi do not consistently express known senescence markers, we sought to characterize the transcripts that are specifically elevated within nevus melanocytes (Fig. 1A). We analyzed previously published datasets comparing RNA expression between benign and malignant melanocytic lesions^26,28^. Each clinical specimen contained matched RNA samples derived from melanocytic nevi and melanomas that arose from those nevi. We reasoned that down-regulated transcripts within these newly-formed melanomas could represent the most immediate barriers to melanocyte growth. Of the differentially expressed genes (adjusted p value <0.05), six of the top ten with elevated expression in nevi were non-coding miRNAs (Fig. 1B and Table S1). Among these were miRNAs previously shown as enriched in nevi compared to melanomas, including MIR211-5p, MIR125A-5p, MIR125B-5p and MIR328-3p^28,29^. To determine whether elevated expression of these miRNAs distinguished nevus melanocytes from normal epidermal melanocytes, we profiled melanocytes from fresh healthy skin and nevi (Table S2). Three of the miRNAs (MIR211-5p, MIR328-3p, MIR125B-5p) were also more highly expressed in melanocytic nevi as compared to healthy melanocytes or melanomas - an expression pattern consistent with enrichment in arrested melanocytes - whereas one miRNA (MIR125A-5p) presented a pattern consistent with lineage specific expression that is lost upon transformation (Fig. 1C). We conclude that MIR211-5p, MIR328-3p, MIR125B-5p are transcripts specifically enriched in arrested melanocytic nevi.

**Figure 1:**
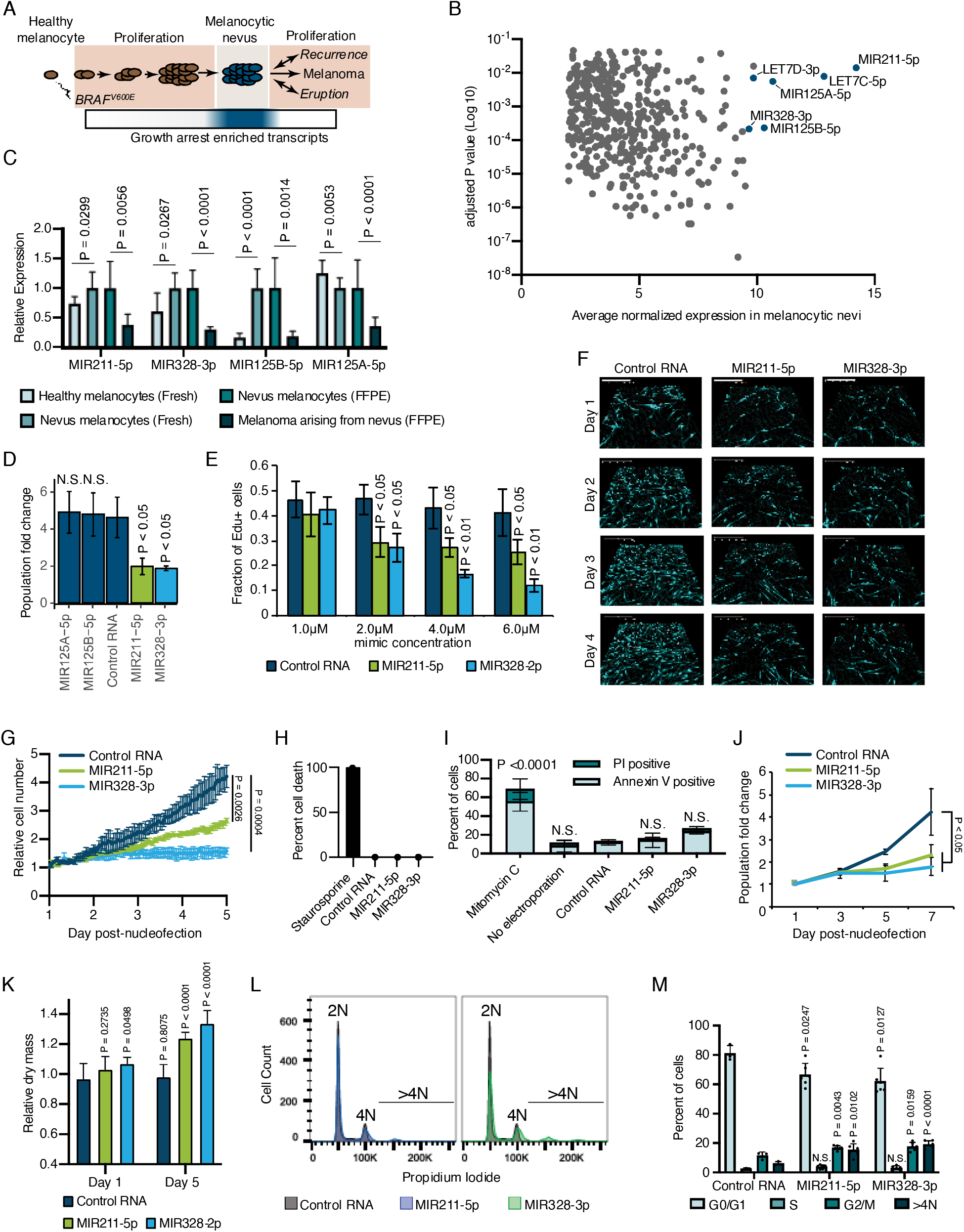
Melanocytic nevus enriched miRNAs induce arrest in primary human melanocytes. **(A)** Schematic of hypothesized plastic nature of BRAF^V600E^- and nevus-associated proliferation arrest. Expression of the predicted nevus enriched transcriptional program is highlighted in blue. **(B)** Scatter plot depicting gene expression combining publicly available sequencing datasets totaling 14 melanoma-arising-from-nevus specimens, dissected into benign and malignant portions (28 total samples). Plotted are the adjusted p values (DESeq2) comparing matched nevus and melanoma samples against average normalized expression in nevus portion. Only genes exhibiting both high expression and significant differential expression in nevus portions are plotted (see Methods). The full list of genes is included as Table S1. **(C)** Left two columns, the relative expression of indicated miRNAs in freshly isolated human melanocytes from BRAF^V600E^ melanocytic nevi (n = 6) compared to BRAF^WT^ healthy skin melanocytes (n = 8). Right two columns, the relative expression of indicated miRNAs in melanocytes dissected from FFPE nevus (n = 7) or melanoma (n = 7) specimens. Data represent mean and standard deviation of normalized RNA sequencing read counts relative to nevus-derived samples. **(D)** Mean and standard deviation for population fold change over seven days of melanocytes nucleofected with indicated miRNA mimics compared to non-targeting control (Control RNA) (n = 3). **(E)** Mean and standard deviation for fraction of EdU positive cells 4 days after nucleofection with indicated concentrations of miRNA mimics (n= 3). P values indicate comparisons to concentration matched control. **(F)** Representative QPI images of melanocytes nucleofected with indicated miRNA mimics. Pixel color indicates low (black), mid (blue) or high (red) optical density. Scale bar = 200µm. **(G)** Mean and standard deviation for relative QPI-derived cell number over time (n = 3). **(H)** Mean and standard deviation for QPI-derived cell death counts as percentage of total cells imaged on day 1 (n = 3). Cells treated with 0.1 µM staurosporine were included as a control (n = 1) for induction of cell death. **(I)** Mean and standard deviation for percent Annexin V/PI positive cells nucleofected with indicated miRNA mimics (n = 3). Cells treated with 5µg/ml mitomycin C were included as a control (n = 3) for induction of apoptosis. **(J)** Mean and standard deviation for relative cell number over time (n = 3). **(K)** Mean and standard deviation for relative QPI-derived dry mass per cell (n = 3). **(L)** Representative histograms of DNA content profiling via propidium iodide incorporation measured by flow cytometry 5 days post-nucleofection of miRNA mimics. **(M)** Mean and standard deviation for percent of cells in indicated phases of cell cycle based upon profiles as in (K) (n = 3-6, individual datapoints shown). P values for D-L calculated by unpaired t tests comparing experimental to Control RNA. N.S. = not significant (P ≥ 0.05)

To investigate whether elevated expression of MIR211-5p, MIR328-3p, or MIR125B-5p induce melanocytic arrest, we introduced synthesized mimetics of the mature miRNAs into primary human melanocytes and tracked proliferation over seven days in culture. The introduction of either MIR211-5p or MIR328-3p resulted in significantly fewer melanocytes compared to a non-targeting control (Fig. 1D). Increased concentrations of these mimics resulted in a dose-dependent decrease in EdU uptake, consistent with a cell cycle defect (Fig. 1E). To better characterize the effect of MIR211-5p or MIR328-3p expression on cell cycle and cell viability, we performed time-lapsed quantitative phase imaging (QPI). We and others have previously established QPI as a method for adherent cell cytometry that can simultaneously measure melanocyte proliferation, cell death, and cell cycle stage with single cell resolution^30–32^. Images were taken every hour for 5 days post-nucleofection (Fig. 1F). Cells expressing MIR211-5p or MIR328-3p exhibited significantly reduced growth curves (Fig. 1G). We observed no evidence of increased cell death (Fig. 1H), and confirmed this result with Annexin V staining (Fig. 1I). Experiments performed using three additional melanocyte preparations counted over seven days post nucleofection yield similar results (Fig. 1J). Thus, MIR211-5p and MIR328-3p induce proliferation arrest in human melanocytes.

Further characterization of the QPI images revealed a distinct increase in intra-cellular dry mass in proliferation-arrested conditions (Fig. 1K). The dry mass of a cell is a measurement of total biomass (DNA, RNA, proteins, lipids, etc) and is quantitatively measured by QPI. Relative dry mass fluctuates with cell cycle stage in a predictable manner^30^. The 20-40% increase we observed here is characteristic of cells that have doubled their DNA content but failed to complete mitosis. We confirmed this finding by measuring DNA content of the arrested populations (Fig. 1L). We observed significant increases of cells with both 4N DNA content and greater than 4N DNA content, indicative of cytokinetic failure following DNA replication (Fig. 1M). Consistent with this interpretation, time-lapsed imaging revealed incidences of tripolar mitosis and anaphase bridging in miRNA-expression melanocytes – both processes associated with cytokinetic failure^33^ - which were not observed in control conditions or standard culture (Video 1 and Figure 1 – figure supplement 1). We conclude that MIR211-5p and MIR328-3p induce mitotic failure and proliferation arrest when introduced into primary human melanocytes.

**Video 1:**
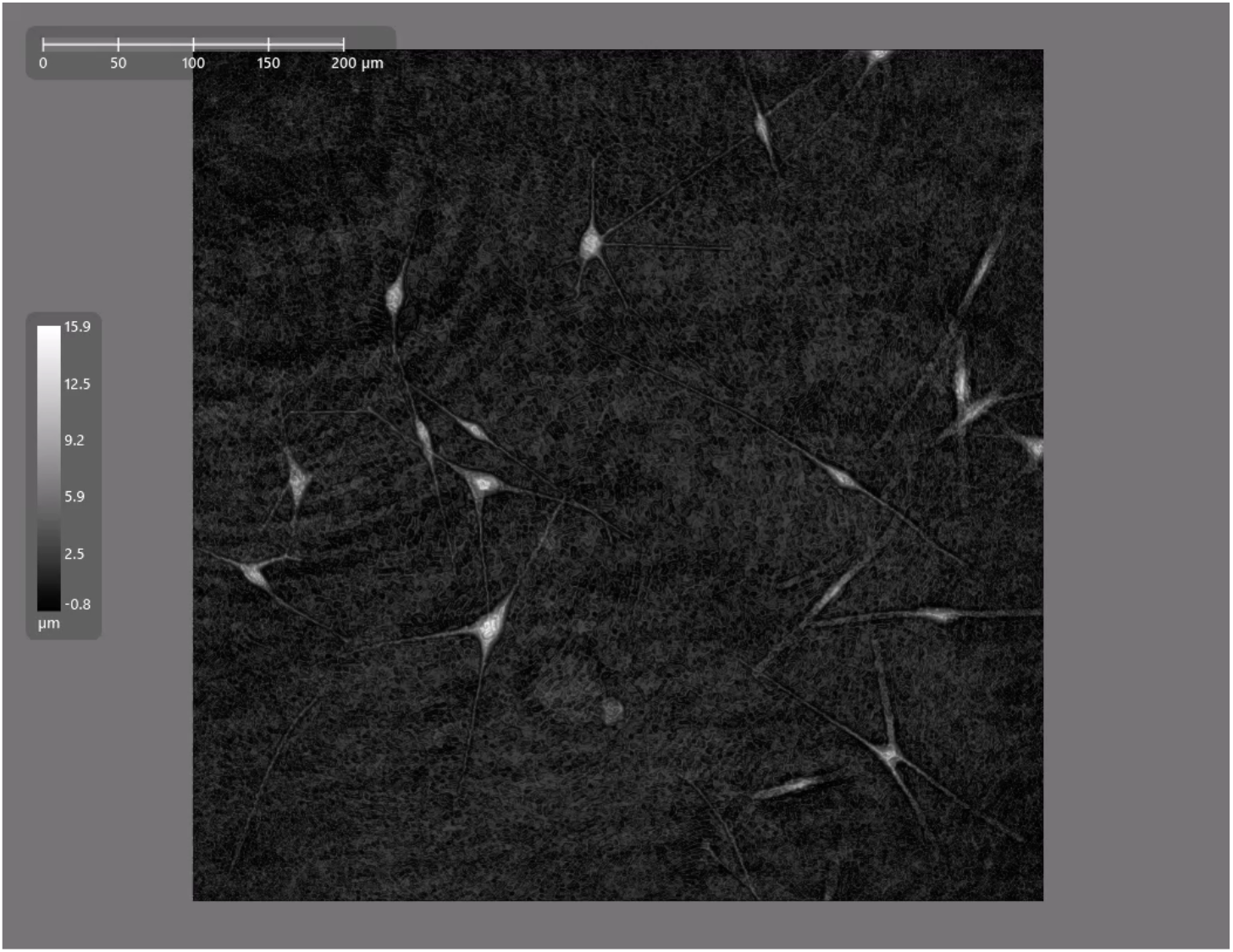
Cytokinesis failure associated with MIR211-5p expression. Four day time-lapse quantitative phase imaging of cytokinesis failure in primary melanocytes transduced with MIR211-5p.

**Figure 1-figure supplement 1.**
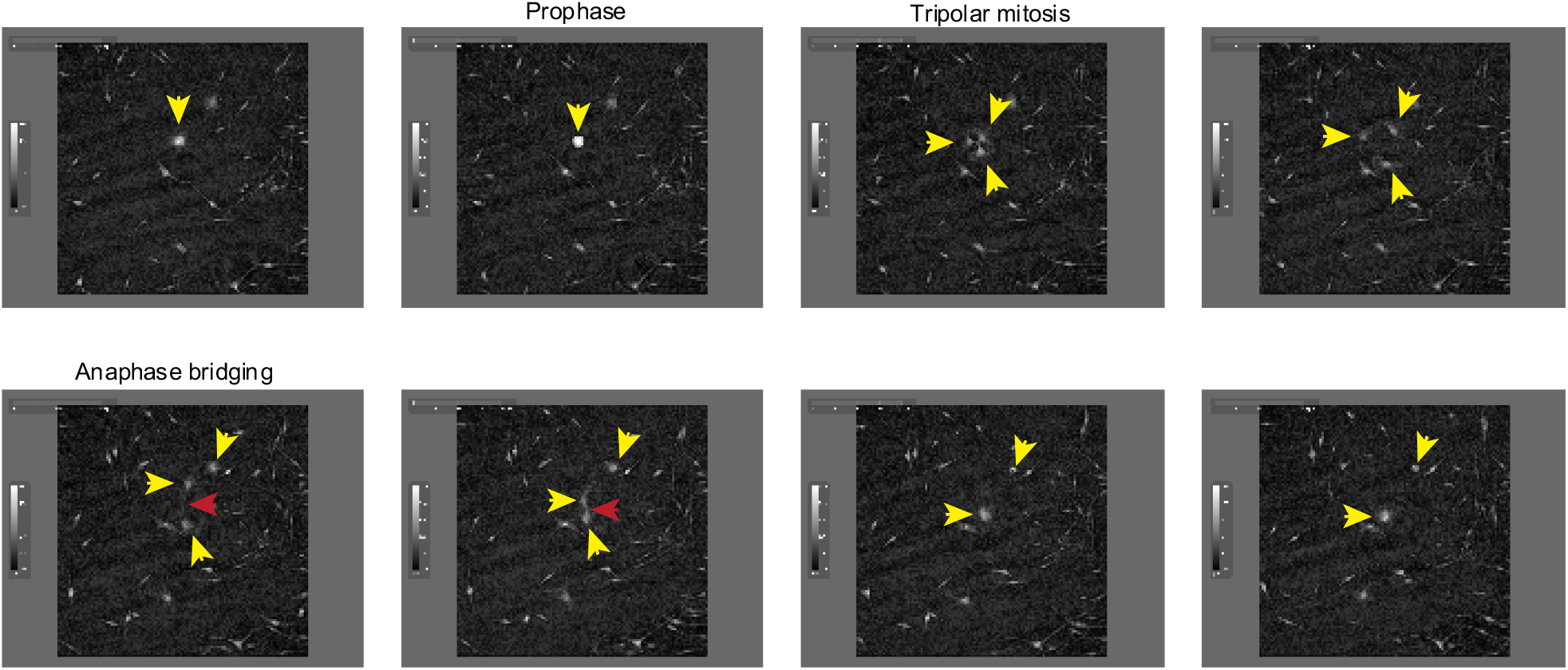
Still frames from Video 1 documenting representative cytokinesis failure in primary melanocytes transduced with MIR211-5p. Yellow arrow heads follow a single cell and its progeny through a tripolar mitosis. Red arrow head follows an anaphase bridge connecting two daughter cells that then remerge.

### Inhibition of AURKB by MIR211-3p and MIR328-5p restricts proliferation in nevi

To investigate the mechanism by which MIR211-5p and MIR328-3p induce arrest in primary human melanocytes, we applied our previously established pipeline for identification of relevant miRNA targets (Fig. 2A)^34^. First, we nucleofected MIR211-5p, MIR328-3p, or control RNA mimics into primary human melanocytes freshly isolated from four independent donors and conducted RNA sequencing (Table S3). We next selected all genes that were significantly downregulated by either miRNA as compared to the control RNA, and further refined the set to the computationally predicted targets of MIR211-5p and/or MIR328-3p. Finally, we nucleofected siRNAs targeting the 115 resulting genes and assayed for reduced EdU incorporation that would phenocopy MIR211-5p or MIR328-3p expression (Table S4). As expected, computationally predicted targets of MIR211-5p and MIR328-3p were enriched in the set of downregulated genes following MIR211-5p or MIR328-3p nucleofection, respectively (Fig. 2B). AURKB, GPR3, and MAPKAPK2 showed the greatest magnitude of repression due to miRNA expression with significant EdU reduction upon siRNA-mediated knockdown (Fig. 2C). Of these, AURKB (predicted MIR211-5p target) and GPR3 (predicted MIR328-3p target), were significantly inhibited upon nucleofection of either miRNA (Fig. 2D). We reasoned these genes might represent convergent nodes driving proliferation arrest. Further exploration of the AURKB transcript revealed a potential MIR328-3p binding site just before the 3’ untranslated region (Fig. 2E). In contrast, we were unable to identify a candidate MIR211-5p binding site in the GPR3 transcript, suggesting indirect inhibition of this gene by MIR211-5p.

**Figure 2:**
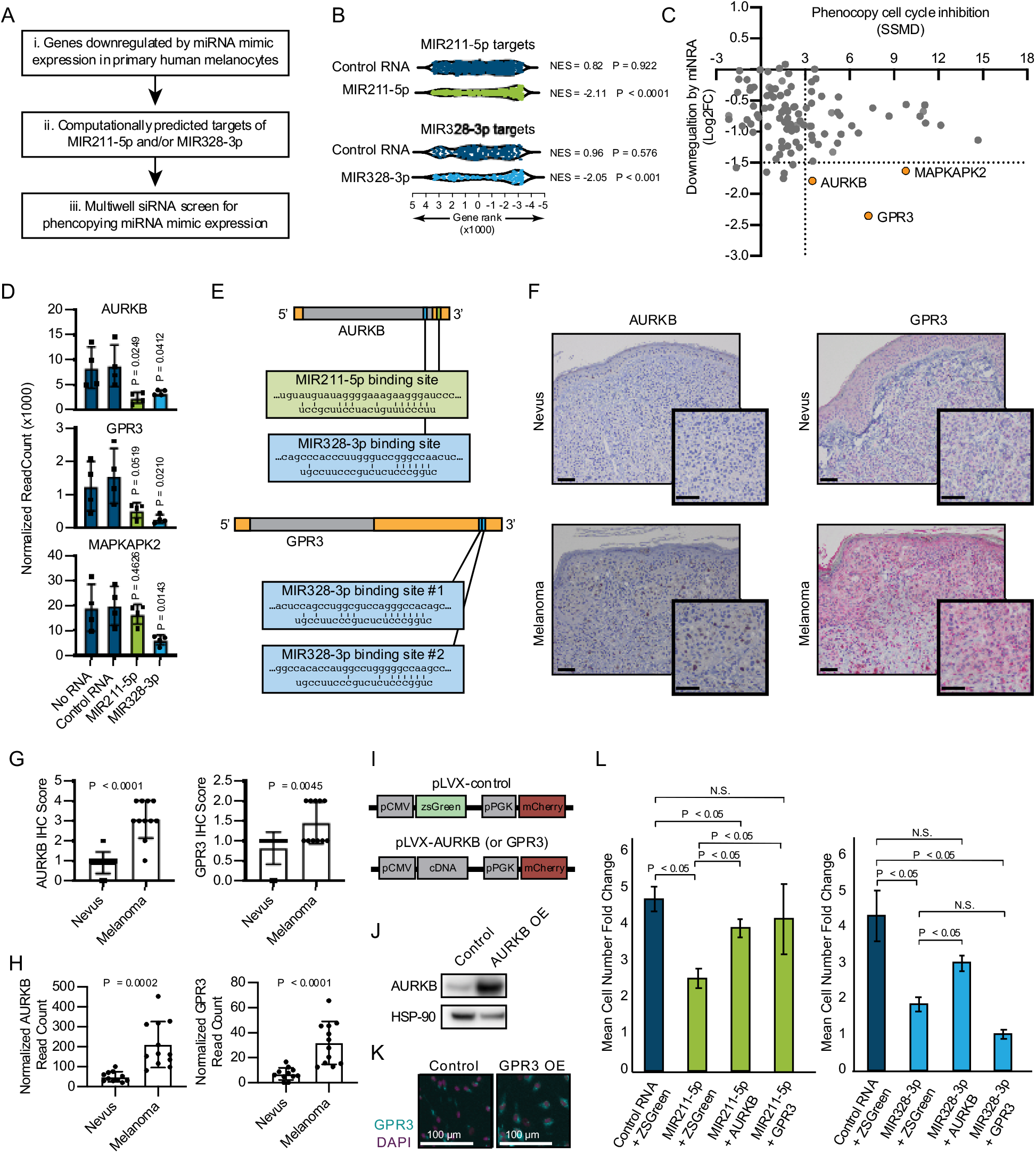
Inhibition of AURKB by MIR211-3p and MIR328-5p restricts proliferation in nevi. **(A)** Schematic of experimental and computational pipeline for identifying targets of MIR211-3p and/or MIR328-3p responsible for proliferation arrest. **(B)** GSEA analysis comparing predicted MIR211-5p and MIR328-3p target mRNAs to changes in gene expression after nucleofection with indicated mimics compared to nucleofection control. NES = normalized enrichment score. P = nominal P value. Table S3 provides full differentially expressed gene lists. **(C)** Plot of siRNA screen results for phenocopying proliferation arrest versus log2 fold change in expression after miRNA mimic nucleofection. Strictly standardized mean difference (SSMD) reports the magnitude and significance of inhibition of Edu incorporation (n = 3, see Table S4 for list of genes and screen results). Dotted lines indicate cut-off values for further evaluation. Orange points indicate genes meeting cut-off criteria. **(D)** Mean and standard deviation for expression of indicated genes after nucleofection of indicated mimics (n = 4). P values calculated by unpaired t tests compared to No RNA. **(E)** Schematics showing predicted binding sites of MIR211-5p and MIR328-3p in AURKB and GPR3 transcripts. **(F)** Representative images of immunohistochemical staining for AURKB (brown chromagen) and GPR3 (red chromagen) expression in FFPE samples of nevi and melanoma. Scale bars = 50µm. **(G)** Mean and standard deviation for immunohistochemical staining scores as in (F) for cohorts of 11 nevi and 11 melanomas. P values calculated by unpaired t tests. **(H)** Mean and standard deviation for read counts for cohorts of 11 nevi and 12 melanomas. P values calculated by unpaired t tests. **(I**) Design of pLVX vectors expressing either AURKB or GPR3. **(J)** Western Blot of AURKB expression in human melanocytes with or without lentiviral AURKB overexpression. HSP90 is the loading control. Source data provided as supplemental files (Fig_2_J_Source). **(K)** Representative photomicrographs (20X) of immunofluorescence for GPR3 (green) or (DAPI) (purple) in human melanocytes with GPR3 lentiviral overexpression (GPR3 OE) or without (Control) over-expression. **(L)** Mean and standard deviation for cell number fold change over 7 days after nucleofection with indicated miRNA mimics in melanocytes transduced with lentiviral constructs expressing zsGreen, AURKB, or GPR3 (n = 6 melanocyte preps, type 2 t test, N.S. = not significant (P ≥ 0.05)). P values calculated by unpaired t tests.

To determine whether decreased AURKB and/or GPR3 expression was associated with melanocytic nevi, we acquired 23 clinical specimens and measured both the transcript and protein expression of these genes (Fig. 2F-H). The expression of both AURKB and GPR3 protein and mRNA were enriched in melanomas as compared to nevi (Fig. 2G-H). Together, these data demonstrate an inverse expression between two miRNAs (MIR211-5p and MIR328-3p) and two targeted mRNAs (AURKB and GPR3) that toggles concurrently with the transition of arrested melanocytic nevi to melanomas. To further investigate whether either mRNA was sufficient to rescue miRNA-mediated arrest, primary melanocytes were transduced with constructs expressing either zsGreen (control), AURKB or GPR3 (Fig. 1I-K) then subsequently nucleofected with either MIR211-5p or MIR328-3p. Expression of either mRNA partially rescued MIR211-5p-induced arrest, while AURKB, but not GPR3, partially rescued MIR328-3p-induced arrest (Fig. 2L). Thus, while other MIR211-5p and MIR328-3p targets, including GPR3, may play a role in the induction of melanocyte proliferation arrest, these data suggest that AURKB inhibition is a critical convergent node of both miRNAs in facilitating proliferation arrest in nevi. This interpretation is further supported by the well-established role of AURKB as an essential orchestrator of successful mitosis^35^.

### BRAF^V600E^ induces a reversible and conditional proliferation arrest in human melanocytes

We hypothesized that AURKB inhibition by MIR211-5p and MIR328-3p may be the underlying mechanism by which BRAF^V600E^ induces proliferation arrest in melanocytes. BRAF^V600E^ expression has been demonstrated to result in an accumulation of growth-arrested fibroblasts with 2N DNA content, presumably in a permanent G0/G1 senescent state^9,12,17^. In contrast, upon overexpression of MIR211-5p or MIR328-3p, we observed an accumulation of arrested melanocytes with ≥4N DNA content, and a significant decrease in cells with 2N DNA content (Fig. 1M). To determine the DNA content profile of primary human melanocytes under BRAF^V600E^-induced arrest, we utilized an established lentiviral construct for dose-responsive doxycycline inducible BRAF^V600E^ expression (Fig. 3A)^25^. We monitored melanocyte growth under a titration of doxycycline and identified the lowest concentration (15.6 ng/mL) that induced sustained arrest after 40 hours exposure (Fig. 3B). Similar to our observations for miRNA-induced arrest, relative per cell dry mass increased by 50% concurrent with BRAF^V600E^ induced-arrest (Fig. 3C). Cell cycle profiling confirmed a significant and dose-dependent accumulation of cells with ≥4N DNA content and a decrease in cells with 2N DNA content (Fig. 3D-E). The cell cycle profiles were notably dissimilar to that of melanocytes treated with a pharmacologic CDK4/6 inhibitor (PD0332991, Palbociclib)^36^ known to induce G0/G1 arrest. Treatment with a pharmacologic AURKB inhibitor (AZD2811, Barasertib)^37^ that induces G2/M arrest and mitotic failure also resulted in an accumulation of melanocytes with ≥4N DNA content and a decrease in cells with 2N DNA content.

**Figure 3:**
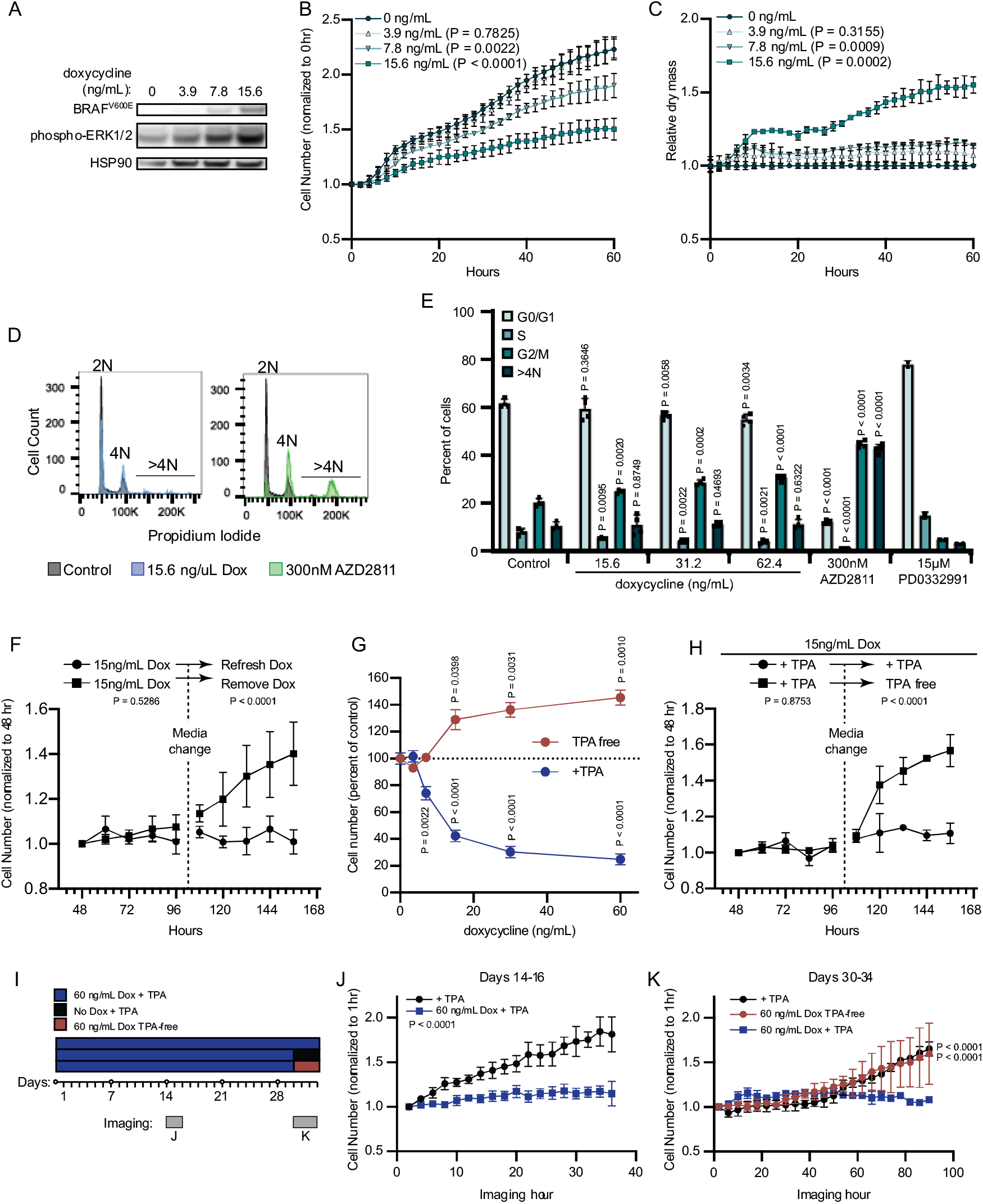
BRAF^V600E^ induces a reversible and conditional arrest in human melanocytes. **(A)** Representative (of n = 3) Western Blot analysis of BRAF^V600E^, phospho-ERK1/2, and HSP90 (loading control) in diBRAF^V600E^ melanocytes in response to increasing concentration of doxycycline. Source data provided in supplemental files (Fig_3A_Source). **(B)** Mean and standard deviation for relative QPI-derived cell number at indicated hours of diBRAF^V600E^ melanocytes in response to increasing concentration of doxycycline (n = 6). P values from unpaired t tests comparing to 0 ng/mL condition. **(C)** Mean and standard deviation for relative QPI-derived dry mass per cell of diBRAF^V600E^ melanocytes in response to increasing concentration of doxycycline at indicated hours (n = 6). P values from unpaired t tests comparing to 0 ng/mL condition. **(D)** Representative histograms of DNA content profiling via propidium iodide incorporation measured by flow cytometry 5 days after treatment with indicated molecules. AZD2811 included as a control for induced cytokinesis failure. **(E)** Mean and standard deviation for percent of cells in indicated phases of cell cycle after treatment with doxycycline based upon profiles as in (D) (n = 4). AZD2811 (n = 4) is a control for induced cytokinesis failure. PD0332991 (n = 2) is a control for G0/G1 arrest. P values from unpaired t tests comparing to Control (no treatment) condition. **(F)** Mean and standard deviation for relative QPI-derived cell number of diBRAF^V600E^ melanocytes at indicated hours after treatment doxycycline. Dotted line indicates change in media to either refresh doxycycline or add media without doxycycline (n = 3). P values from unpaired t tests comparing final timepoints. **(G)** Mean and standard deviation for cell number as percent of control of diBRAF^V600E^ melanocytes cultured with or without TPA after 5 days of treatment with indicated concentrations of doxycycline. P values from unpaired t tests comparing to 0 ng/mL conditions. Baseline population doubling times were 3.2 days (+TPA) and 4.2 days (TPA-free). **(H)** Mean and standard deviation for relative QPI-derived cell number of diBRAF^V600E^ melanocytes at indicated hours after treatment doxycycline. Dotted line indicates change in media to either refresh TPA-containing media or add media without TPA (n = 3). P values from unpaired t tests comparing final timepoints. **(I)** Schematic of experimental design testing the reversibility of growth arrest after 30 days exposure to high doxycycline (60 ng/mL). Days of live imaging represented in (J) and (K) are indicated by grey bars. **(J)** Mean and standard deviation for relative QPI-derived cell number of diBRAF^V600E^ melanocytes at 14 days after addition of doxycycline (n = 3). P values from unpaired t tests comparing final timepoints. **(K)** Mean and standard deviation for relative QPI-derived cell number of diBRAF^V600E^ melanocytes at 30 days after addition of doxycycline (n = 3). Media was changed just prior to imaging to either refresh doxycycline (blue), +TPA media without doxycycline (black) or add TPA-free media with doxycycline (red). P values from unpaired t tests comparing final timepoints.

In addition to the accumulation of 2N cells, previous reports of RAF expression in fibroblasts show that transient expression can induce a durable arrest phenotype^17^. To test the durability of BRAF^V600E^-induced arrest in human melanocytes, we induced doxycycline-dependent BRAF^V600E^ expression for 48 hours and monitored cell number for an additional two days to confirm arrest. Media was then replaced to remove doxycycline. Within twelve hours of doxycycline removal, the previously arrested melanocytes began to proliferate (Fig. 3F), demonstrating that BRAF^V600E^-induced proliferation arrest in melanocytes is reversible and dependent on continued oncogene expression.

Nevi recurrence after excision^18^ and eruption upon external stimuli^19^, together with our observation that BRAF^V600E^ induced arrest is reversible in melanocytes, suggested that BRAF^V600E^-induced arrest may depend on secondary signals from the microenvironment. Tetradecanoylphorbol acetate (TPA) is a protein kinase C (PKC) activator commonly used as a mitogen in primary melanocyte media to substitute for endothelin receptor type B activation^38^. In contrast, TPA also inhibits the cell cycle of melanoma cells^39,40^. BRAF^V600E^ expression permits growth of the Melan-A cell line in the absence of TPA^41^. We therefore wanted to determine the relative contributions of TPA and BRAF^V600E^ to the BRAF^V600E^-induced primary melanocyte arrest *in vitro*. We induced BRAF^V600E^ expression in primary human melanocytes in two medias – one containing TPA (+TPA) or one containing the endothelin receptor type B ligand, endothelin-1 (TPA-free) and observed divergent dose-responsive phenotypes (Fig. 3G). In the presence of TPA, BRAF^V600E^ expression inhibited cell proliferation. By contrast, in TPA-free media, BRAF^V600E^ expression induced proliferation. We next subjected melanocytes to the arrest-inducing conditions of doxycycline treatment in +TPA media for four days, then replaced with doxycycline-containing, TPA-free media. We observed a rapid shift to the proliferative phenotype even in the continued presence of doxycycline (Fig. 3H). To determine whether the observed reversibility of BRAF^V600E^-induced growth arrest was due to insufficient duration or degree of expression of the oncogene, we exposed melanocytes to a high concentration of doxycycline (60ng/mL) for 30 days prior to switching medias (Fig. 3I). Melanocytes in +TPA media remained arrested for the duration of doxycycline treatment then grew readily upon removal of either doxycycline or TPA (Fig. 3J-K). From these results, we conclude that BRAF^V600E^ expression induces divergent and interconvertible phenotypes – either hyperproliferation or a reversible arrest – in human melanocytes conditional on extrinsic stimuli.

### BRAF^V600E^-induced arrest is dependent on melanocyte growth conditions and differentiation state

To investigate the relative contributions of BRAF^V600E^ and TPA to the expression of MIR211-5p and MIR328-3p, we conducted small RNA sequencing of melanocytes grown in +TPA or TPA-free media and with or without BRAF^V600E^ expression. The expression of MIR328-3p was relatively stable, demonstrating less then 2-fold changes across conditions (Fig. 4A-B, Table S5). In contrast, MIR211-5p was one of the most differentially expressed genes in +TPA conditions compared to TPA-free conditions (adj. P = 6.8E-23) (Fig. 4A-B, Table S5). Induction of BRAF^V600E^ in +TPA conditions did not further alter expression of either miRNA. Direct comparison of the arrested melanocytes in +TPA +Dox conditions to the proliferative melanocytes in -TPA +Dox conditions revealed MIR211-5p as the most significantly downregulated miRNA in the proliferative condition (Fig. 4B). Thus, in comparison to TPA, BRAF^V600E^ induction had minimal effect on the expression of the miRNAs. The most notable effect was a decrease of MIR328-3p expression coinciding with the induction of BRAF^V600E^ in TPA free conditions (Fig. 4A). While MIR328-3p downregulation likely contributes to increased proliferation in this condition, the more pronounced effect of TPA on MIR211-5p expression suggests that the mechanisms governing whether BRAF^V600E^ induces arrest or proliferation are a function of the cell’s transcriptional state rather than directly downstream of BRAF^V600E^ itself.

**Figure 4:**
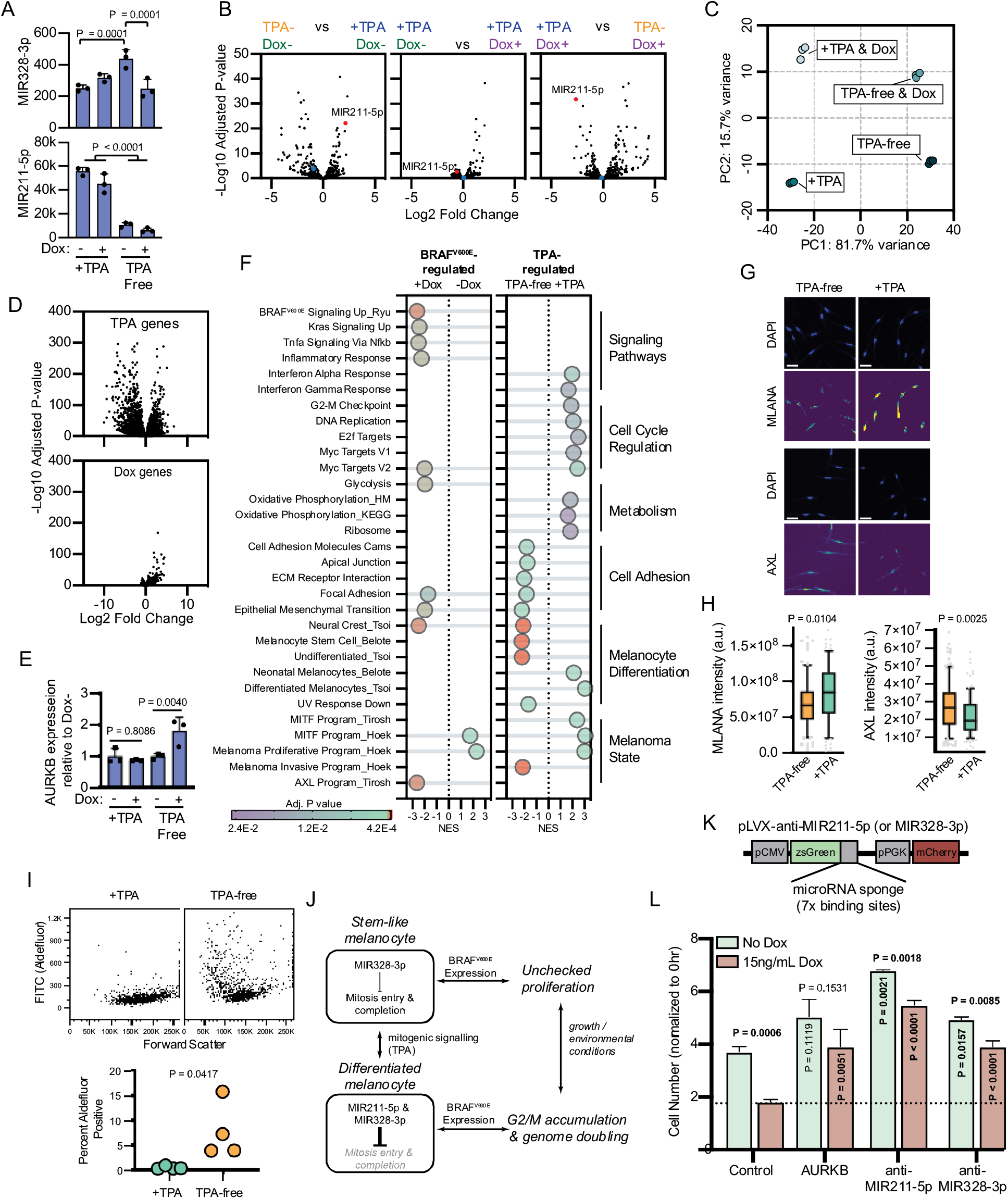
BRAF^V600E^-induced arrest is dependent on melanocyte growth conditions and differentiation state. **(A)** Normalized read counts for MIR328-3p (top) and MIR211-5p (bottom) in TPA-containing (+TPA) or TPA-free media, with or without 15.6 ng/mL doxycycline (Dox). See Table S5 for individual values and exact P values. **(B)** Volcano plots depicting log2 fold change against adjusted P value (-Log10) from differential expression (DE) analysis of small RNA sequencing of indicated comparisons. MIR211-5p labelled and shown in red. MIR328-3p shown in blue. **(C)** First and second principal components (PC) separating diBRAF^V600E^ melanocytes cultured in indicated conditions. **(D)** Volcano plots depicting log2 fold change against adjusted P value (-Log10) from differential expression (DE) analysis of mRNA sequencing of all +TPA specimens versus all TPA-free specimens (top) or all Dox specimens versus all no-Dox specimens (bottom). **(E)** Normalized read counts for AURKB normalized to no Dox conditions (n = 3). P values from unpaired t tests. Sequencing data represented in A-F are from n=3 per condition analyzed with DESeq2. **(F)** Gene set enrichment analysis (GSEA) comparing DE gene sets from (D) to Molecular Signature Database Hallmark gene sets, KEGG Pathway gene sets and published signatures of melanocyte signaling and differentiation from Belote, 2021; Tsoi, 2018; Tirosh, 2016; Ryu, 2011; and Hoek, 2010. All enrichments with adjusted P value < 0.025 are depicted. See Table S6 for all associations. Normalized enrichment score (NES). Enrichment differences between pathways across the four conditions in (C) are depicted in Figure 4-figure supplement 1. **(G)** Representative photomicrographs of immunofluorescence for MLANA or AXL in human melanocytes cultured in +TPA or TPA-free conditions. White scale bar = 30µm. **(H)** Quantification of fluorescence intensity (arbitrary pixel intensity units, a.u.) from experiments represented in (G). Box (median, 25^th^ and 75^th^ percentiles) and whiskers (10^th^ and 90^th^ percentiles) of 84, 88, 113, and 96 cells respectively. P values from unpaired t tests. **(I)** Representative plots (top) and quantification (bottom, n = 4) of relative aldehyde dehydrogenase activity of melanocytes grown in +TPA or TPA-free conditions. **(J)** Schematic of model supported by transcriptomic analyses. TPA induces a more differentiated state in human melanocytes. Due to increased MIR211-5p and base-line MIR328-3p expression, AURKB expression is capped, resulting in G2/M accumulation upon BRAF^V600E^ expression. **(K)** Design of pLVX vectors expressing microRNA sponges fused to zsGreen. **(L)** Mean and standard deviation for cell number as percent of time point 0 hr after 7 days of culture in +TPA media with or without 15.6 ng/mL Dox (n = 3 per condition). Melanocytes were transduced with either pLVX-control, -AURKB, -anti-MIR211-5p, or -anti-MIR328-3p. P values from unpaired t tests comparing to matched no Dox condition (horizontal), Control no Dox condition (vertical in green bar) or Control Dox condition (vertical in red bar).

To better characterize the two transcriptional states influencing the BRAF^V600E^ phenotype, we conducted mRNA sequencing of the same specimens. Principal component (PC) analysis revealed that 97.4% of the variance between samples was explained by two principal components – PC1 (81.7% variance) separated transcriptomes based upon exposure to TPA and PC2 (15.7% variance) separated transcriptomes based upon BRAF^V600E^ expression (Fig. 4C, Table S5). We conducted differential expression analyses to compare all samples exposed to +TPA versus TPA-free media, regardless of BRAF^V600E^ expression (Fig. 4D, top) and to compare all samples expressing BRAF^V600E^, regardless of TPA exposure (Fig. 4D, bottom). As expected from the PC analysis and similar to the miRNA profiles, most gene expression variance was due to media conditions with a smaller, but consistent, set of genes activated by BRAF^V600E^.

One initially surprising result of this analysis was the static expression of both MIR211-5p and MIR328-3p in +TPA conditions, regardless of BRAF^V600E^ expression. However, as inhibitors of translation, miRNA function depends on the expression level of its targets^42,43^, effectively functioning as buffers of gene expression that prevent aberrantly high levels of a transcript^44^. BRAF^V600E^ has been previously demonstrated as an upstream activator of AURKB transcription in melanoma cells^45^. We therefore reasoned that MIR211-5p expression in +TPA conditions might dampen AURKB activation by BRAF^V600E^. Consistent with this hypothesis, the addition of doxycycline induced AURKB expression in TPA-free / MIR211-5p low conditions, but this effect was not observed in +TPA / MIR211-5p high conditions (Fig. 4E).

We next conducted gene set enrichment analyses (GSEA) comparing TPA-regulated genes and BRAF^V600E^-regulated genes to Molecular Signature Database Hallmark gene sets^46^., the KEGG PATHWAY database^47^, and signatures important in BRAF^V600E^ signaling^48^, human melanocyte differentiation^49^ or melanoma cell state^50–52^. Among the pathways uniquely enriched with BRAF^V600E^ expression were the expected increases in signatures downstream of BRAF^V600E^, MAPK (KRAS) signaling, and glycolysis^53^ (Fig. 4F and Table S6). Gene sets uniquely enriched in +TPA media included signatures associated with cell cycle regulation, such as the G2-M checkpoint, oxidative phosphorylation, the MITF program, and differentiated melanocytes. In contrast, TPA-free media induced gene sets associated with cell adhesion and melanocyte progenitor and stem cells. Corroborating this observation, melanocytes grown in TPA-free media expressed higher levels of AXL (Fig. 4G-H), lower levels of MLANA, a melanocyte differentiation marker and presented greater activity of aldehyde dehydrogenase, enzymatic activity associated with stemness (Fig. 4I). These results indicate that exposure to TPA induces a more differentiated melanocyte phenotype, consistent with previous reports^54–56^.

To look for synergist effects of both BRAF^V600E^ expression and media conditions, we performed pair-wise differential expression analyses of each condition (Table S6 and summarized in Figure 4-figure supplement 1). Generally, most gene sets identified in Figure 4F were regulated by only one of the two stimuli. For example, genes downstream of BRAF^V600E^ signaling were only influenced by doxycycline exposure. Interesting exceptions to this trend were the inflammatory response, which required both stimuli; cell cycle regulation gene sets, which were activated by TPA alone but further elevated when supplemented with doxycycline; and epithelial to mesenchymal transition genes, which were activated by either stimulus. A subset of melanocyte dedifferentiation gene sets, specifically neural crest genes, the melanoma invasion program and the AXL program, but not markers of melanocyte stem cells, were activated by either TPA-free media or BRAF^V600E^ expression, albeit the activation was more pronounced in the former. This observation suggests that BRAF^V600E^ expression alone does induce some programs associated with melanocyte dedifferentiation, but with the retention of MIR211-5p expression, the induction is insufficient to overcome growth arrest.

Collectively, our transcriptomic analyses support a model whereby growth in +TPA conditions induces a more differentiated state in melanocytes coinciding with an increase in MIR211-5p expression (Fig. 4J). This increased MIR211-5p combined with relatively stable MIR328-3p expression, dampens entry into mitosis and/or successful completion of mitosis at least in part through AURKB inhibition. The attenuation of cell division results in an accumulation of cells in G2/M – an effect that is more pronounced in conditions stimulating accelerated growth, such as with BRAF^V600E^ expression. To test this model, we transduced doxycycline-inducible BRAF^V600E^ melanocytes in +TPA conditions with lentiviral constructs constitutively expressing either AURKB (Fig. 2I-J) or miRNA sponges (Fig. 4K) to either MIR211-5p or MIR328-3p. AURKB expression had no baseline effect on melanocyte growth, but rescued BRAF^V600E^-induced arrest (Fig. 4L), suggesting the inability of the oncogene to induce AURKB in this context serves as a bottleneck in cell division. Also consistent with the model, sponges against either MIR211-5p or MIR328-3p increased baseline division rate and rescued BRAF^V600E^-induced arrest, demonstrating that both miRNAs contribute to inhibition of melanocyte division, with affects that are more pronounced in the presence of growth signals.

**Figure 4 - figure supplement 1.**
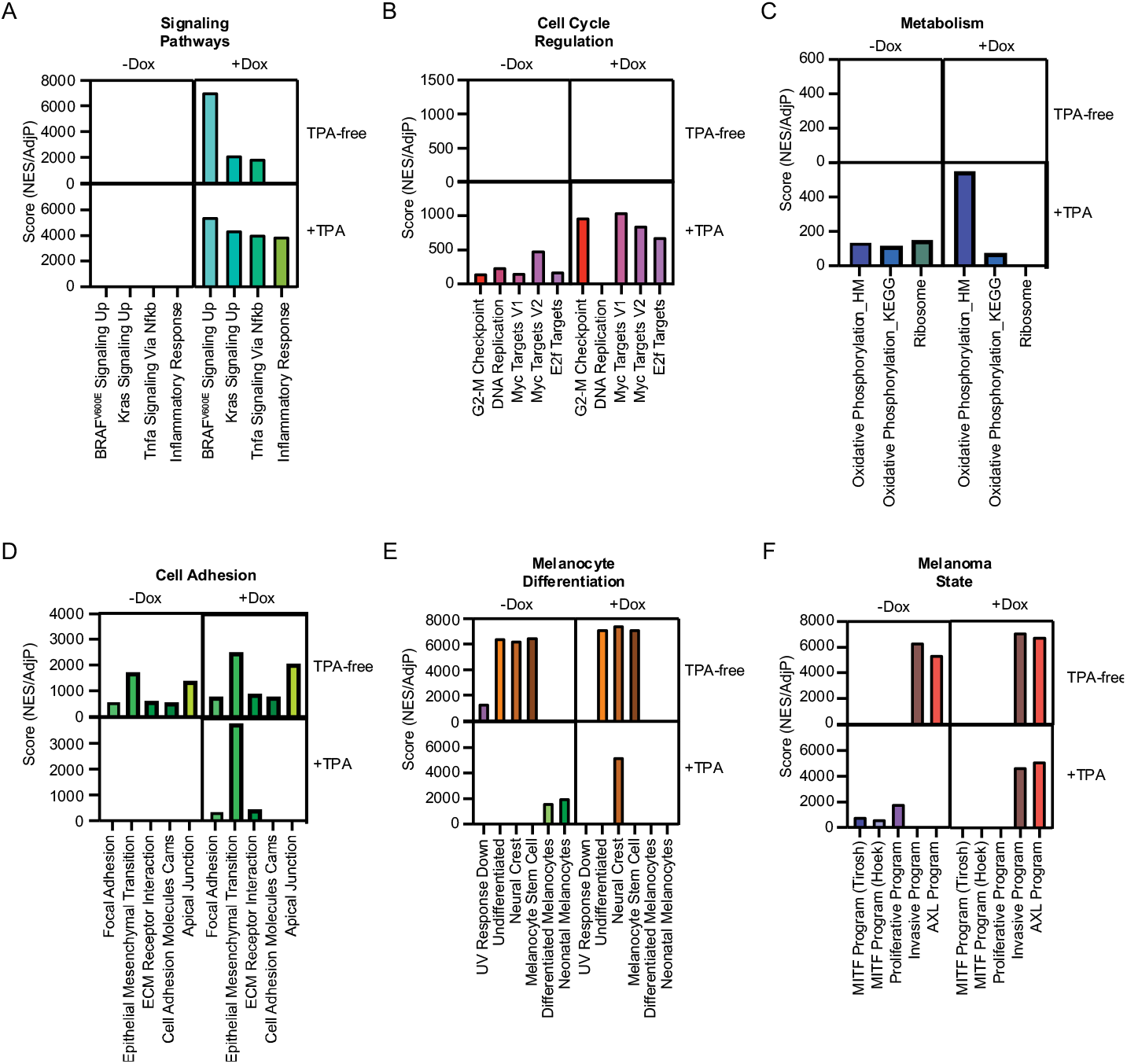
Depiction of gene set enrichment for gene sets in Figure 1F across the four conditions in Figure 4C. Pair-wise comparisons were conducted as in Figure 1F, comparing the +TPA -Dox condition to other conditions. Scores were generated by dividing positive NES values by the AdjP value. For the +TPA -Dox quadrant, scores were averaged when occurring in more than one pair-wise comparison.

### AURKB inhibition drives melanocytic nevus-associated proliferation arrest

To further investigate whether this mechanism of arrest occurs in clinical nevus specimens, we examined an independent cohort of 15 nevi for evidence of mitotic failure. Histopathologic analysis revealed that all nevi within the cohort exhibited binucleation (Fig. 5A). The percent observed binucleated melanocytes ranged from 3-12% (mean = 6.3%). It is important to note that since binucleation can only be observed in the fraction of cells sectioned perpendicular to the plane of the adjacent nuclei, 3-12% is likely an under-estimate. We also observed giant multinucleated cells in 10 of the 15 nevi, indicative of multiple rounds of DNA synthesis and cytokinetic failure and consistent with previous reports^57^.

**Figure 5:**
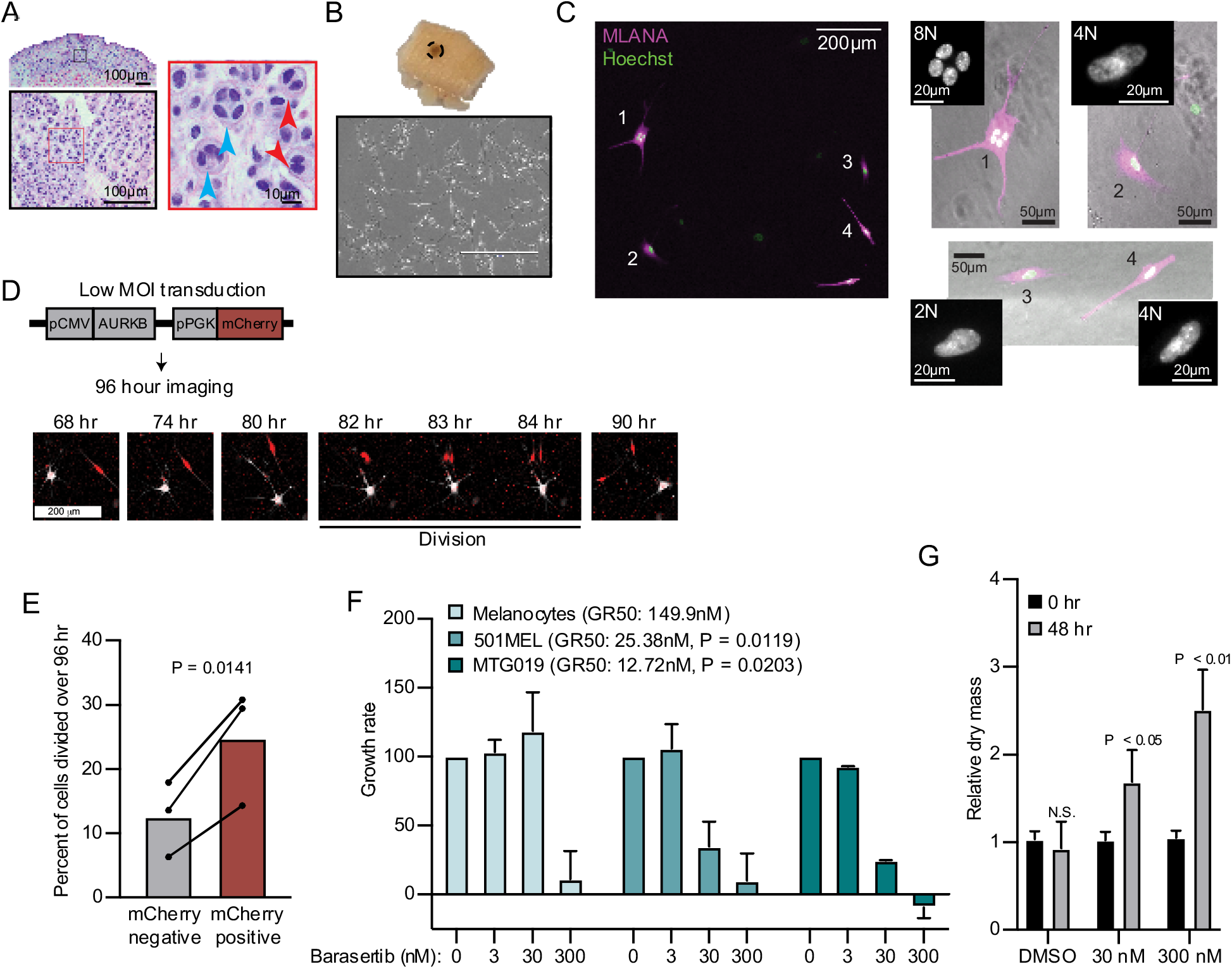
AURKB inhibition is critical for BRAF^V600E^-induced arrest of human nevi. **(A)** Representative image of 15 H&E-stained melanocytic nevus. Border colors indicate two consecutive magnifications. Arrowheads indicate bi- (red) or multi- (blue) nucleation. **(B)** Example images of skin specimen containing a melanocytic nevus (circled) and phase contrast microscopy of melanocytes isolated from nevus portion. Scale bar = 400µm. **(C)** Images of MLANA (purple) and Hoeschst (green) co-staining of melanocytic nevus derived melanocytes in culture. Zoomed images show representative 2N, 4N, and 8N cells. **(D)** Representative of images of QPI coupled with fluorescence microscopy to identify adjacent mCherry -expressing and mCherry-negative melanocytes. Individual cells were identified at time zero, tracked and monitored for division for 96 hrs. **(E)** Mean and individual matched datapoints for the percent of n = 3 mCherry-negative or mCherry-positive cells identified at time zero that divided over 96 hrs. P value from paired t test. **(F)** Mean and standard deviation for 48 hr growth rates of indicated cells treated with Barasertib. P values from unpaired t tests comparing the 30nM samples of melanoma lines to primary melanocytes (n = 3). **(G)** Mean and standard deviation for relative QPI-derived dry mass per 501MEL cell at initial time point (black) compared to 48 hr treatment with Barasertib (grey) (n = 3). P values from unpaired t tests.

We sought to determine whether AURKB inhibition was necessary and sufficient for the multi-nucleated arrest we observed in melanocytic nevi. We obtained three fresh skin biopsies containing nevi, microdissected the nevus portion, and isolated melanocytes (Fig. 5B). Nevus-derived melanocytes appeared healthy in culture. Consistent with the histopathologic cohort, co-staining with melanocyte marker, MLANA, and Hoechst revealed frequent 4N and 8N melanocytes (Fig. 5C). Transduction of nevus melanocytes with an AURKB and mCherry expressing lentiviral vector at low multiplicity of infection (MOI) showed that the mCherry positive cells divided significantly more than the their mCherry negative counterparts (Fig. 5D-E) in cells from all three nevus donors. This result suggests that expression of AURKB is also rate limiting for the proliferation of human nevus melanocytes, as we observed in primary melanocytes expressing BRAF^V600E^.

To test the sufficiency of AURKB inhibition to limit BRAF^V600E^ hyperproliferation in melanoma, we assessed normalized growth rate inhibition of three cultures - primary human melanocytes, an established BRAF^V600E^ human melanoma cell line, and a short-term BRAF^V600E^ human melanoma culture - when exposed to a pharmacologic AURKB inhibitor (Barasertib). Normalized growth rate measurements (such as GR_50_) are independent of baseline cell division rate and are demonstrably more reproducible than the commonly calculated half maximal inhibitor concentration (IC_50_) when comparing different cell cultures^58^. An additional advantage of GR_50_ is the ability to distinguish growth rate inhibition (GR_50_ approaches 0) versus cell death (GR_50_ is negative). Since GR_50_ calculations require proliferation curves, we conducted live QPI over 48 hours with a range of Barasertib concentrations (0-300nM). Primary melanocytes were largely resistant to the compound with reduced growth apparent only at supraphysiological levels (Fig. 5F). In contrast, both melanoma cultures responded to ten-fold lower dosage of the compound with GR_50_ approaching zero, indicating primarily arrest, not death. In these conditions, the per cell dry mass also increased significantly, consistent with a G2/M arrest (Fig. 5G). Taken together, these data demonstrate that AURKB inhibition limits the rate of BRAF^V600E^-induced proliferation in human melanocytic nevi and melanomas.

## Discussion

Cellular senescence is conventionally defined as a permanent cell-cycle arrest associated with a variety of molecular markers and an irreversible arrest in the G0/G1 phase of the cell cycle^10–12^. The vast majority of melanocytic nevi are stably arrested and harbor BRAF^V600E^, an oncogenic mutation that induces cellular senescence when expressed in a variety of mammalian cells^9,12,17^. While these observations seemingly implicate OIS as the mechanism driving nevus-associated proliferation arrest, a substantial body of evidence has challenged this paradigm in recent years^8,9,18–25^. Here, we provide further evidence to challenge the model that acquisition of BRAF^V600E^ induces premature cellular senescence in melanocytes and uncover an alternative mechanism driving nevus formation and stability. We propose that acquisition of the BRAF^V600E^ mutation does not necessitate growth arrest, but instead permits melanocytes to toggle between hyperproliferation and mitotic arrest. Our transcriptomic analyses demonstrate that external stimuli can modulate the differentiation state of BRAF^V600E^ melanocytes, orchestrate the expression level of MIR211-5p, and inform which phenotype is triggered by the oncogene – before or after the oncogene is introduced.

The senescence model is largely rooted in seminal work demonstrating that sustained ectopic expression of BRAF^V600E^ induces proliferative arrest in primary human melanocytes^9^ - an experiment reproduced by several independent labs^24,25,59,60^, including our own^23^. In each of these studies, melanocytes were cultured in TPA containing media. Thus, while observations presented here – that BRAF^V600E^ -induced proliferative arrest is reversible and conditional on the presence of TPA – provide new insights that challenge the senescence model, they do not contradict the data presented in these previous reports.

One inherent limitation of using any *in vitro* approach to investigate the mechanisms underlaying clinical phenomena is an inherent incongruity in the magnitude of duration. Melanocytic nevi frequently persist for several decades, whereas *in vitro* experiments are generally conducted on the order of days to weeks. In previous *in vitro* studies, the periods of time melanocytes were monitored subsequent to BRAF^V600E^ introduction ranged from one to three weeks. In studies utilizing an inducible RAF in human fibroblasts, 24 hours of expression was sufficient to drive a durable proliferative arrest, even upon removal of the oncogene^17^. In this study, we induced growth arrest with high levels of BRAF^V600E^ for thirty days and observed the arrest was still reversible. It remains possible that an even longer exposure to BRAF^V600E^ would elicit a permanent proliferative arrest consistent with senescence. It is noteworthy, however, that the identified mechanisms driving proliferative arrest *in vitro* were also supported by analyses of clinical nevi specimens.

While supported by analyses of both primary human melanocytes and clinical specimens, one limitation to this report is the lack of *in vivo* confirmation using animal models. One well-established genetically engineered mouse model of melanoma expresses BRAF^V600E^ from the endogenous locus in tyrosinase expressing cells with spatiotemporal control. The predominant phenotype of BRAF^V600E^ activation in this model is melanocytic hyperplasia, which increases in prominence with BRAF^V600E^ dose^6^. While growth-arrested melanocytic “nevus-like” lesions also form in this model, they are not associated with markers of senescence^8^. More recently, an elegant study utilizing a zebrafish model of melanoma demonstrated that driving BRAF^V600E^ expression from promoters associated with melanocyte progenitors, but not differentiated melanocytes, resulted in melanoma initiation^61^. These observations are all consistent with our data and support the conclusion that BRAF^V600E^-induced proliferation in melanocytes is dependent on differentiation state.

The presented model offers an explanation for several clinical phenomena. First, melanocytic nevi form when BRAF^V600E^ drives hyperproliferation followed by proliferation arrest. Second, nevus eruption involves expansion of previously stable nevi over a short period of time. Third, incomplete excision of melanocytic nevi can result in subsequent regrowth of the benign lesion^18^. Each event requires modulation of the BRAF^V600E^ melanocyte between a proliferative and arrested phase. Our data demonstrate that BRAF^V600E^ melanocytes can indeed oscillate between proliferative and arrested states and are consistent with each of these phenomena. Our interpretation assumes that once acquired, BRAF^V600E^ expression is constant and that the environmental context of the melanocytes is variable, serving as a “gatekeeper” to BRAF^V600E^-induced proliferation. However, if BRAF^V600E^ expression also fluctuates, then an equally plausible interpretation of our data is that BRAF^V600E^ serves as a “gatekeeper” to environmentally induced proliferation. Interestingly, nevus eruption has been observed in metastatic melanoma patients treated with single agent selective BRAF inhibitors^62–66^, consistent with our observation that proliferation arrest in melanocytes requires sustained expression of the oncogene. A more thorough understanding of the responsible environmental stimuli in physiologic conditions, discussed in more detail below, would help to clarify which of the mechanisms – variable BRAF^V600E^, variable environment, or both – underlie nevus formation, eruption and recurrence.

It is important to note that while our data support a model of reversible proliferation arrest in nevi, we do not suggest this reversal is commonplace. Unquestionably, the vast majority of melanocytic nevi present with sustained arrest. However, we do propose that the duration of arrest is dynamic and modulated by exposure to external signals. Since our data suggest that loss of MIR211-5p and MIR328-3p expression and restoration of AURKB expression is concurrent with progression to melanoma, identification of the source of cell-extrinsic signals might suggest new strategies for chemoprevention or therapy. We have identified TPA, a known activator of PKC signaling, as one extrinsic factor that induces expression of MIR211-5p and a more dedifferentiated state. Previous studies have identified TPA as either a mitogen, inhibitor of the G1/S transition, or inhibitor of the G2/M transition, dependent on the context^39,40,67–69^. We demonstrate that *in vitro*, TPA activation of MIR211-5p is required for BRAF^V600E^-induced mitotic failure. Yet, the inclusion of TPA in primary melanocyte culture is artificial and the environmental stimulus that regulates MIR211-5p expression in skin remains unknown. One hypothesis is that PKC activation from keratinocyte interactions serves as the extrinsic stimulus. MIR211-5p is reported to be transcriptionally regulated by both MITF^70^ and UV exposure^71^. However, TPA-free media containing an endothelin receptor type B ligand, endothelin-1, did not elicit MIR211-5p expression or BRAF^V600E^ arrest, suggestive that the downstream signaling pathways regulated by the two molecules are not identical. PKC activation can also result in increased MAPK signaling^72,73^ raising the possibility that internal MAPK fluctuations may play a role in toggling between BRAF^V600E^-induced proliferation versus arrest. The term “mitogenic window” describes the specific range of oncogene doses that elicit a hyperproliferative phenotype. The mitogenic window for MAPK signaling activation can be remarkably fine. Here, we conducted a two-fold dilution series of doxycycline but observed no evidence of a mitogenic window. However, it remains possible that we simply missed the “sweet spot” in TPA conditions, and that a finer dilution series would reveal a mitogenic window even in the presence of TPA. Regardless, our data demonstrate that in TPA conditions the mitogenic window for BRAF^V600E^ expression is, at least, exceedingly narrow, whereas in non-TPA conditions the window is wide open. An important future direction will be to identify the nature and physiological source of this signal for melanocytic nevi in human skin.

Diminished MIR211-5p/MIR328-3p expression concurrent with increased AURKB expression occurs in a substantial portion of melanomas. AURKB expression was previously reported as significantly elevated in melanoma metastases^45^ and here we report increased expression in primary melanomas. Similarly, MIR211-5p expression is consistently observed as lost in melanoma as compared to melanocytic nevi across independent studies^27–29^. Indeed, MIR211-5p expression levels classify nevi from melanomas with a high degree of accuracy^27,28^. We have reported on the diagnostic potential of MIR328-3p expression as well, although multi-study validation was impeded by the absence of hybridization probes in earlier microarray-based profiling datasets^28^. *In vitro*, we have now demonstrated that inhibition of either miRNA is sufficient to block BRAF^V600E^-induced arrest in melanocytes. These data suggest that mitotic failure serves as a barrier to melanoma progression and is consistent with the genome duplication and copy number alterations associated with invasive melanoma^4,74^. In melanoma, the specific cause of this genomic instability is unknown, but reentry of cells into cell cycle after mitotic failure has been shown to cause genome duplication and accumulation of copy number alterations in other cancers^75^. Further experiments will be necessary to determine whether dysregulation of the spindle checkpoint in nevi contributes directly to the copy number variations present in melanomas.

In summary, we have identified that BRAF^V600E^ induces a reversible arrest in human melanocytes orchestrated by MIR211-5p/MIR328-3p regulation of AURKB and conditional on the melanocyte differentiation state. We present a mechanistic model that allows for melanocytic nevus formation, eruption and recurrence. Moreover, our observations provide an explanation for the genetic duplication and instability inherent to invasive melanomas and open new directions for novel chemopreventative and therapeutic strategies.

## Competing Interests

R.L.J and M.L.W are inventors on international patent application no. PCT/US2019/023834 concerning the use of microRNAs investigated in this article as molecular diagnostics for melanoma.

**Acknowledgments**

This work was supported by the National Institutes of Health (DP5OD019787 and R01CA229896 to R.L.J., 5F31CA236377 to A.S.M., P30CA042014 to University of Utah), the University of California, San Francisco Program for Breakthrough Biomedical Research (PBBR) Sandler Fellowship to R.L.J. and the 5 for the Fight fellowship to R.L.J. We acknowledge the use of the HCI Shared Resources for Research Informatics (RI), Cancer Biostatistics (CB), High-Throughput Genomics and Bioinformatics Analysis (GBA), and the Biorepository and Molecular Pathology (BMP) Research Histology Section supported by P30CA042014 awarded to HCI from the National Cancer Institute.

## Author Contributions

A.S.M: Investigation, Formal analysis, Writing – original draft, Visualization, Funding acquisition; R.L.B.: Methodology, Investigation, Resources, Writing – review & editing; H.Z.: Methodology, Investigation, Validation, Writing – review & editing; M.U.: Investigation; K.B.: Investigation; R.T.: Formal analysis; M.C. – Formal analysis; A.H.S.: Data curation, Writing – review & editing; R.A., Writing – review & editing: Resources; S.H.: Conceptualization; D.H.L.: Conceptualization, Writing – review & editing; T.H.M.: Resources; M.W.V.: Conceptualization; D.G.: Resources, Writing – review & editing; M.W.: Resources, Writing – review & editing; U.E.L.: Investigation, Formal Analysis, Resources, Writing – review & editing; R.L.J.: Conceptualization, Formal analysis, Investigation, Writing – original draft; Visualization, Supervision, Funding acquisition.

## Materials and Methods

### Transcriptomic profiling

For FFPE samples, histopathologic review, microdissection, targeted exon sequencing, phylogenetic analysis, and RNA and small RNA sequencing of each tumor area were previously described (phs001550.v2.p1)^23,26,28^. For this study, small RNA reads were aligned to human reference (hg37) with Bowtie^76^ and small RNA reference groups (miRBase21) were counted. For mRNA sequencing, reads were aligned to human reference (hg37) with Bowtie2 and counted with HTSeq. Differential expression analysis was performed using DESeq2^77^ with p-values adjusted by Benjamini-Hochberg method (p-adj). Previously published RNA sequencing datasets from *Shain et al. 2018* and *Torres, et al*. 2020 were re-analyzed here with DESeq2 the combined for visualization. For RNA profiling from primary melanocytes, total RNA was extracted using TRIzol Reagent (Thermo Fisher, 15596-026) and further purified with the RNeasy Power Clean Pro Cleanup Kit (Qiagen, 13997-50) to remove melanin. For small RNAseq of nevus melanocytes sequencing libraries were constructed with the TailorMix Small RNA Library Preparation Kit (SeqMatic, CA) and sequencing was performed on the Illumina HiSeq2500 platform at single-end 50bp. For small RNAseq of cultured BRAF^V600E^ transduced melanocytes, sequencing libraries were constructed with the Qiagen QIAseq miRNA Library Prep kit and sequencing was performed on the NovaSeq SP platform at paired-end 50bp. For mRNAseq of microRNA mimic nucleofected melanocytes, 150bp paired end sequencing was conducted by GeneWiz. For mRNAseq of cultured BRAF^V600E^ transduced melanocytes, sequencing libraries were constructed with the Illumina TruSeq Stranded mRNA Library Prep kit and sequencing was performed on the NovaSeq S4 platform at paired-end 150bp.

### Cell derivation and culture

Benign human melanocytic nevi were excised with informed consent from patient donors at the UCSF Dermatology clinic (San Francisco, CA) or HCI Dermatology clinic according to Institutional Review Board-approved protocols. Nevus tissue was minced and enzymatically dissociated in 1mg/mL collagenase (Sigma 11213857001) and 3.3mg/mL dispase (Sigma, 4942078001) in DMEM (Thermo Fisher, 10569044) for 1 hour at 37°C. Melanocytes were further isolated by 5-day exposure to 10µg/mL G418 (InvivoGen, ant-gn-1). BRAF status was confirmed via sanger sequencing (Quintarabio) using the primer set: BRAF forward: 5’-GCA CGA CAG ACT GCA CAG GG -3’; BRAF reverse: 5’-AGC GGG CCA GCA GCT CAA TAG -3’. BRAF wildtype (normal) human melanocytes were isolated from de-identified and IRB consented neonatal foreskins or adult skin. Foreskin tissue was incubated overnight at 4°C in dispase and epithelia were mechanically separated from the dermis the following morning. Epithelial tissue was minced and incubated in 0.25% trypsin (Gibco 25200056) for 4min at 37°C. Trypsin was quenched and tissue centrifuged at 500xg for 5min at room temperature. The cell/tissue pellet was resuspended in Melanocyte medium (ThermoFisher, M254500) with HMGS (ThermoFisher, S0025) and plated in low volume.

The 501Mel human melanoma line (Gift from Dr. Boris Bastian, CVCL_4633) were cultured in RPMI with 10% Fetal Bovine Serum (FBS, VWR, S107G), 1% Penicillin-Streptomycin (Gibco, 15140122), 1% L-Glutamine (Gibco, 25030149), and 1% Non-Essential Amino Acids (Gibco, 11140050). HCIMel019 was derived from patient-derived xenograft (PDX) tumors propagated in mice. An HCIMel019 (P2) subcutaneous tumor was resected from mouse, minced in digestion buffer (100μM HEPES (Gibco, 15630-080), 5% FBS (DENVILLE, FB5001-H), 20μg/ml gentamicin (Gibco, 15710-064), 1x insulin (Gibco, 51500-056), and 1mg/ml collagenase IV (Gibco, 17104-019) in DMEM (Gibco, 11965-092)), and digested overnight at 37°C. Cells were filtered through a 100µm filter (Falcon, 352360) and red blood cells removed with RBC lysis buffer (0.5M EDTA, 0.5M KHCO3 (Sigma, 237205), and 5M NH4CL (Sigma, A9434)). Remaining cells were washed with PBS (Gibco, 10010023), and cultured at 37°C and 5% CO2 in Mel2 media consisting of 80%MCDB153 (Sigma, M7403), 20% L15 (Gibco, 11415-064), 2% FBS (DENVILLE, FB5001-H), 1.68mM CaCL (Sigma, C4901), 1x insulin (Gibco, 51500-056) 5ng/ml EGF (Sigma Aldrich, E9644), 15μg/ml Bovine Pituitary Extract (Gibco 13028-014), and 1x Pen/strep (Gibco, 15070-063).

### Small RNA Nucleofection

Human melanocytes were trypsinized (Gibco, 25300062), quenched, and centrifuged at 300xg. Cells were resuspended in R Buffer (ThermoFisher, MPK1025) at 10,000 cells/µL. 10µL of cell slurry was mixed with miRIDIAN microRNA mimics (Dharmacon: hsa-MIR211-5p C-300566-03-0005, hsa-MIR328-3p C-300695-03-0005, MIRControl 1 CN-001000-01-05 or MIRControl 2 CN-002000-01-05); or On-TargetPlus siRNA (Dharmacon, custom library); at 4µM final concentration and nucleofected using the NEON Transfection System and protocol (Thermo Fisher, MPK5000).

### Generation of Lentiviral vectors

pTRIPZ-diBRAFV600E was a gift from Todd Ridky^25^. pLVX-AURKB (Addgene #153316) and pLVX-GPR3 (Addgene #153317) were generated by subcloning the respective human cDNA (from Addgene #100142 and #66350) into the MluI and BamHI sites] of the pLVX-Che-hi3 vector (a gift of Sanford Simon)^78^. pLVX-anti-MIR211-5p (Addgene #153318), pLVX-anti-MIR328-3p (Addgene #153319), and pLVX-Che-zsGreen (Addgene #153320) were generated by inserting zsGreen with or without a 3’UTR into the MluI and XbaI sites of the pLVX-Che-hi3 vector. The 3’UTRs contained 7 tandem microRNA binding sites to report and inhibit microRNA function, as previously described^34^. ZsGreen was subcloned from pHIV-zsGreen (gift from Bryan Welm & Zena Werb (Addgene #18121))^79^.

### Lentiviral transduction

2.75×10^6^ HEK293T cells were plated on 10cm tissue culture dishes and grown for ∼24 hours in DMEM with 10% FBS. For each 10cm plate, 5µg lentiviral vector, 3.3µg of pMLV-GagPol and 1.7µg of pVSV-G packaging plasmids were added to 500µL of jetPRIME buffer and 20µL of jetPRIME transfection reagent (Polyplus, 712-60), and transfected according to manufacturer’s instructions. 48 hours post-transfection, viral supernatant was collected and filtered using 0.45µm syringe filters (Argos 4395-91). Human melanocytes seeded at 1.0×10^5^ – 2.0×10^5^ cells/well density in 6-well plates were incubated in viral supernatant with 10µg/mL polybrene (Sigma, TR-1003) and centrifuged at 300xg for 60 min at room temperature. The viral supernatant was removed and replaced with growth media. Transduced cells were either selected with puromycin (1µg/mL for 5 days, Sigma P8833) or sorted for mCherry expression using a BD FACSAria II. Cells expressing pTRIPZ-diBRAFV600E were treated with doxycycline (Sigma, D9891) at indicated concentrations.

### Live Quantitative Phase Imaging

Live quantitative imaging was performed using either the HoloMonitor M4 imaging cytometers (Phase Holographic Imaging, Lund, Sweden) or the Livecyte platform (Phasefocus, Sheffield, UK). Analyses of cell proliferation, dry cell mass, and death were acquired with the M4 platform and analyzed using HStudio (v2.6.3) as previously described^31^. For each experiment, human melanocytes were seeded into 6-well plates (Sarstedt, 83.3920) at 100,000 cells/well and either live-imaged or serially imaged as indicated. Analysis of growth rate were acquired with the M4 platform and analyzed using App Suite (v3.2.0.60) as previously described^58^. For each experiment, 100,000 melanocytes, 60,000 501Mel cells or 150,000 HCIMel019 cells were plated per well and media containing indicated concentrations of barasertib (AURKB inhibitor; AZD1152-HQPA | AZD2811, Selleckchem, A1147) was added. Cells were imaged for 48-72 hours. Analyses of fluorescent reporters coupled with quantitative phase imaging was conducted using the Livecyte platform and analyzed using Analyze (v3.1) (Phasefocus, Sheffield, UK).

### EdU assays

Normal human melanocytes were nucleofected with microRNA mimics, LNA microRNA inhibitors, or infected with lentiviral vectors (as above) and seeded in 48-well plates at a density of 50,000 cells/well. 4 days post-seeding EdU was added to culture media at a 10µM final concentration. 24 hours after EdU addition, media was removed and cells were stained for EdU incorporation and nuclei using the Click-iT EdU Imaging Kit (Thermo Fisher, C10337) and protocol. Images of EdU and nuclei staining were acquired using the Evos FL microscope and quantified with FIJI.

### Western Blotting

Protein was collected using RIPA Buffer (Thermo Fisher, 89901) supplemented with HALT Protease Inhibitor (Thermo Fisher, 87786). Immunoblotting was carried out as previously described^23^. Membranes were incubated overnight at 4 °C with primary antibodies at the following dilutions: anti-HSP90 (CST, 4874) 1:1000, anti-BRAFV600E (Spring Bioscience Corp, E19292) 1:1000, anti-phosphoERK1/2 (CST, 4970) 1:1000, anti-AURKB (Abcam, ab2254) 1:1000. Membranes were washed 4X with TBS and 0.5% Tween20, incubated with HRP-conjugated secondary antibody (1:2000) for 30 minutes at room temperature, and visualized with Lumina Forte Western HRP substrate (Millipore, WBLUF0500).

### Flow analyses

For assessing DNA content, trypsinized melanocytes (as above) were resuspended in 400 PBS. 1mL -20 °C 200 proof ethanol was added dropwise while gently vortexing to achieve 70% final concentration for fixation and incubated overnight at -30 °C. Cells were centrifuged at 800xg for 5 min, washed with fresh PBS, and incubated in 500µL FxCycle PI/RNase Staining Solution (Invitrogen, F10797) for 20 min at 37 °C. Data were collected on the Fortessa (BD) at low speed and analyzed with FlowJo v10.7.1. Annexin V/PI staining was performed by trypsinizing melanocytes as above. Cells were washed with PBS and resuspended with Annexin V binding buffer (BioLegend, 422201) at a concentration of 1×10^6 cell/mL. 5µL APC Annexin V antibody (BioLegend, 640919) and 10µL PI were added. Cells were gently vortexed and incubated in the dark at room temperature for 15 min. Cells were resuspended in 400µL of Annexin V Binding Buffer. Data were collected on the FACs Verse (BD) and analyzed with FlowJo v10.7.1.

Aldehyde dehydrogenase activity was measured first by trypsinizing melanocytes as above and then using the ALDEFLUOR kit (Stemcell Technologies, 01700) as described in the manufacture’s protocol.

### Human nevus and melanoma tissue immunohistochemistry

The archived clinical specimens used in the study were procured with IRB approval from the UCSF Dermatopathology Pathology Service archives. For assessment of AURKB and GPR3 expression, 11 melanoma and 11 melanocytic nevi with previous diagnosis were de-identified and re-verified histologically (UEL). For assessment of DNA content, 15 additional melanocytic nevi with previous diagnosis were re-verified histologically (UEL). Tissue was fixed in 10% neutral-buffered formalin, processed, embedded in paraffin, and stained with hematoxylin and eosin. 4µm FFPE sections were stained with AURKB (1:400 dilution, Abcam ab2254) or GPR3 (1:125 dilution, Abnova H00002827-M01) antibodies. UEL reviewed immunohistochemical stains with semiquantitative grading for GPR3 (0=none; 1=patchy positive; 2=strong positive) and AURKB (0=none; 1=rare positive; 2=scattered positive; 3= frequent positive; 4=many positive). For in vitro immunofluorescence, cells were fixed and stained as previously described^80^ with AXL (1:250 dilution, Cell Signaling #8661) or MLANA (1:100, Abcam ab731).

### Statistical analyses

For differential gene expression, p values were calculated with the DESeq2 (v1.30.1) default Wald test adjusted by the Benjamini-Hochberg method using a 5% false discovery rate (FDR)^81^. Pathway analyses were analyzed using the fast gene set enrichment package^82^ in R with a 10% FDR. Gene set enrichment analyses were conducted using GSEA (v4.0.3, Broad Institute) Preranked tool with 1000 permutations. For *in vitro* experiments, pilot studies were initially conducted in triplicate. The required minimal sample size to assess the difference between independent means with independent standard deviations with an alpha error probability of 0.05 and a power of 0.95 was calculated using G*Power (v. 3.1.9.4). P-values were calculated using either paired or unpaired two-tailed T Tests or Wilcoxon tests via Prism 8 (Graphpad) as indicated in figure legends.

### Experimental set-up

For cohorts of clinical specimens, sample number (n) refers to the number of specimens, each from a different patient. For *in vitro* experiments, n refers to independent experiments conducted on different days. In cases where fresh primary melanocytes were used, the cells in each experiment are derived from an independent donor. In cases where cell lines are used, the cells in each experiment represent a different passage and/or day of that line. For different conditions within experiments, replicate wells were plated and arbitrarily chosen for each condition (e.g. not treated versus treated; different concentrations of a compounds, etc.). All data were included.

## Ethics statement

The archived clinical specimens used in the study were procured with IRB approval from the UCSF Dermatopathology Pathology Service archives (11-07951). Fresh human tissue samples used in this study were obtained with informed consent and consent to publish from each patient. Samples were first coded before being provided to the researchers. The study protocol conformed to the ethical guidelines of the Declaration of Helsinki (1975). Samples were processed following standard operating procedures approved by IRB at University of California, San Francisco (16-20299) and at the Huntsman Cancer Institute (00124195).

## Data availability

The data which support these findings are available within the article and its supplementary files, as well as from the corresponding author upon reasonable request. The sequencing data produced in this manuscript is available from GEO under the accession number GSE150849.

## Supplemental Table Legends

**Supplemental Table S1 (related to Figure 1b)**

Results of differential expression analyses performed on previously published databases of matched melanocytic nevi and melanoma-arising-from-nevi.

**Supplemental Table S2 (related to Figure 1c)**

Normalized read counts from small RNA sequencing performed on melanocytes derived from healthy human skin and melanocytic nevi.

**Supplemental Table S3 (related to Figure 2)**

Results of differential expression analyses performed on primary melanocytes nucleofected with microRNA or control mimics.

**Supplemental Table S4 (related to Figure 2c)**

Results from siRNA screen for EdU incorporation performed in primary melanocytes. Mean and standard deviation of triplicate screens shown for each gene. Column D indicates the microRNA predicted to targeted the gene.

**Table S5 (related to Figure 4)**

Results of differential expression analyses performed on primary melanocytes with or without BRAF^V600E^ in +TPA or TPA-free conditions.

**Table S6 (related to Figure 4)**

Results of GSEA analyses performed on primary melanocytes with or without BRAF^V600E^ in +TPA or TPA-free conditions.

**Fig_3A_Source and Fig_2J_Source**

Zip files containing raw and annotated images of western blots.

