## Supplementary figures and images for "BRAF^V600E^ induces reversible mitotic arrest in human melanocytes via microRNA-mediated suppression of AURKB"

### figure_3A_source_data_BRAFv600e_annotated.tiff

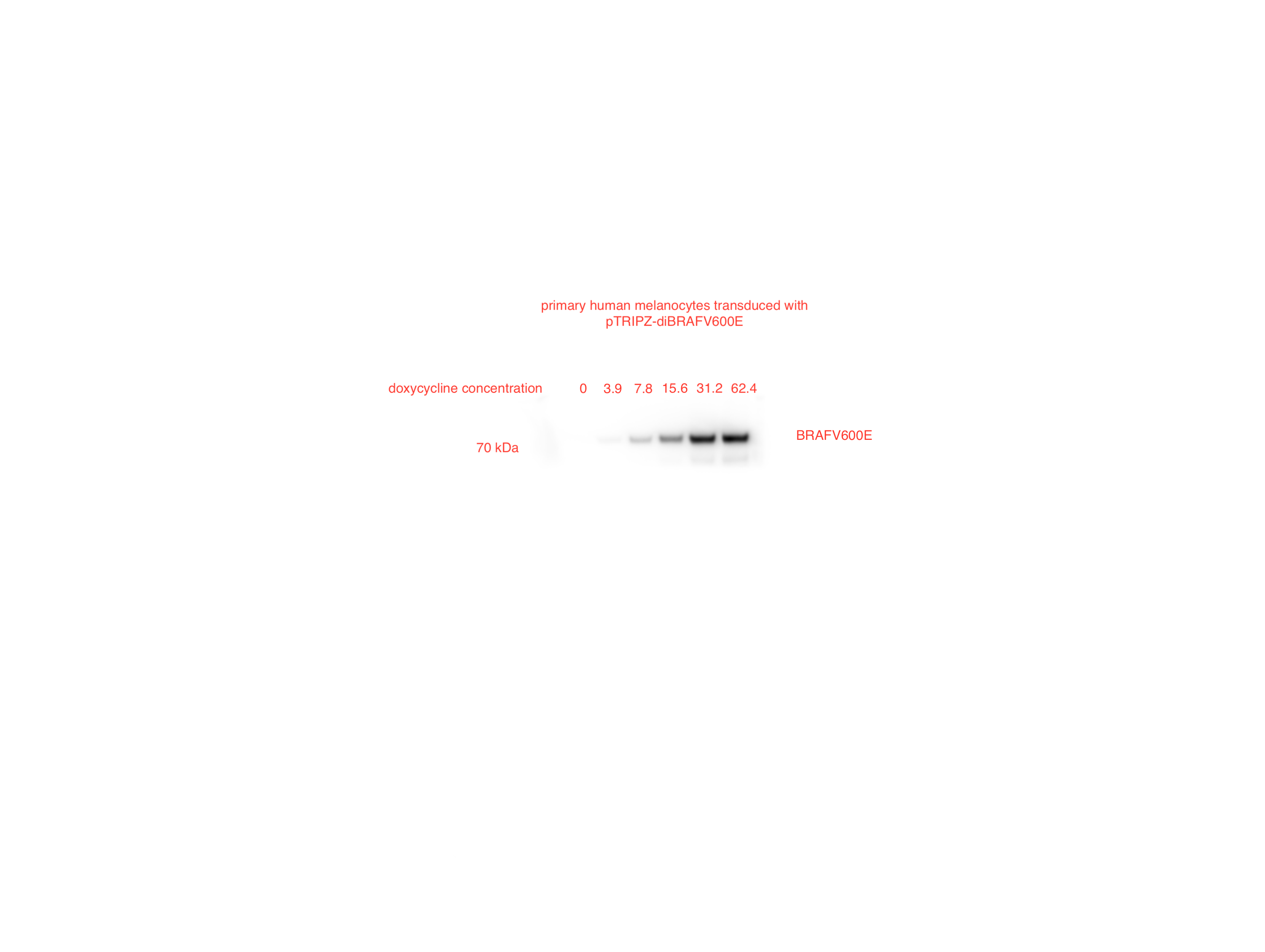

### figure_3A_source_data_BRAFv600e_pub.tif

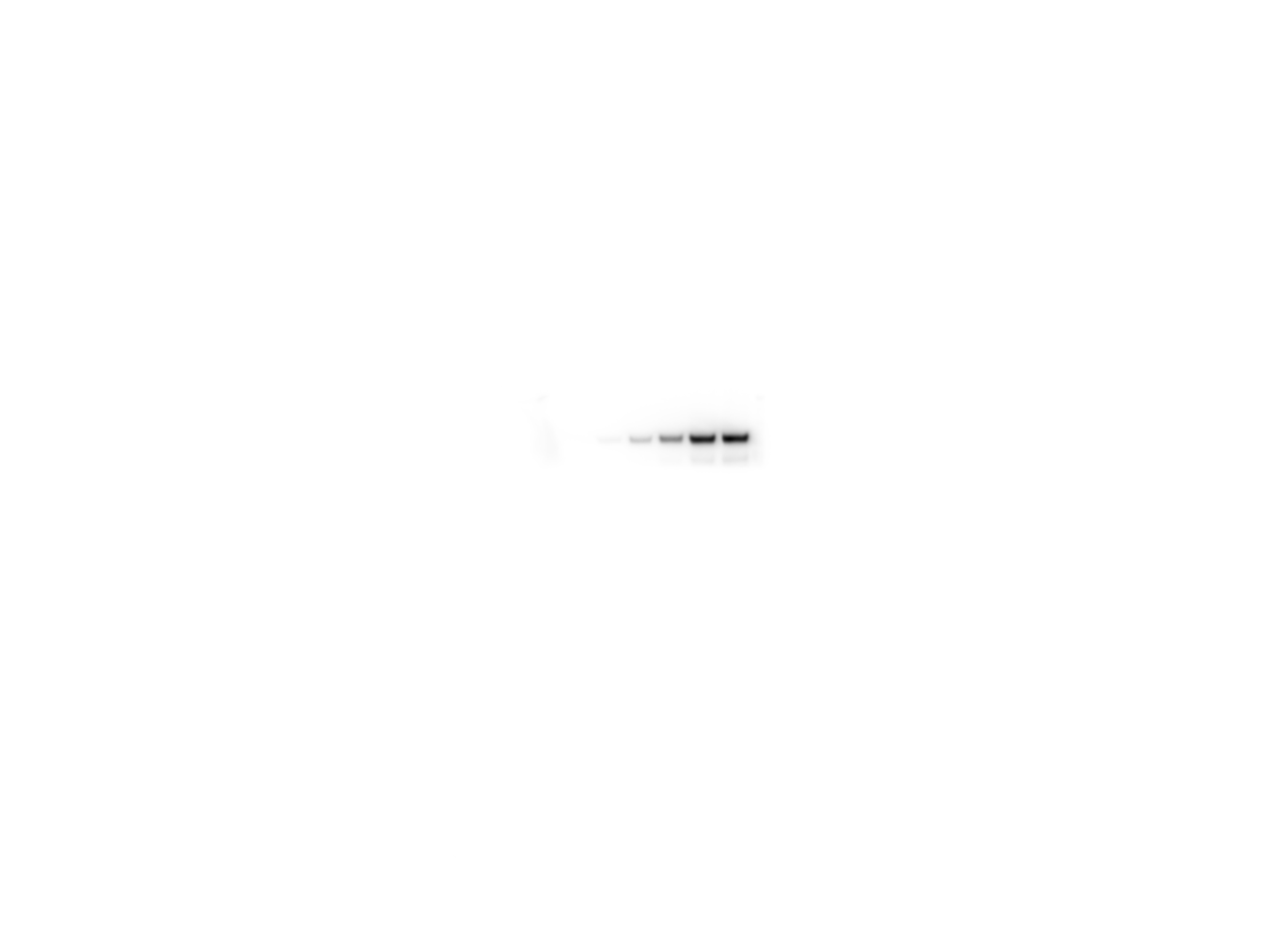

### figure_3A_source_data_hsp90.tif

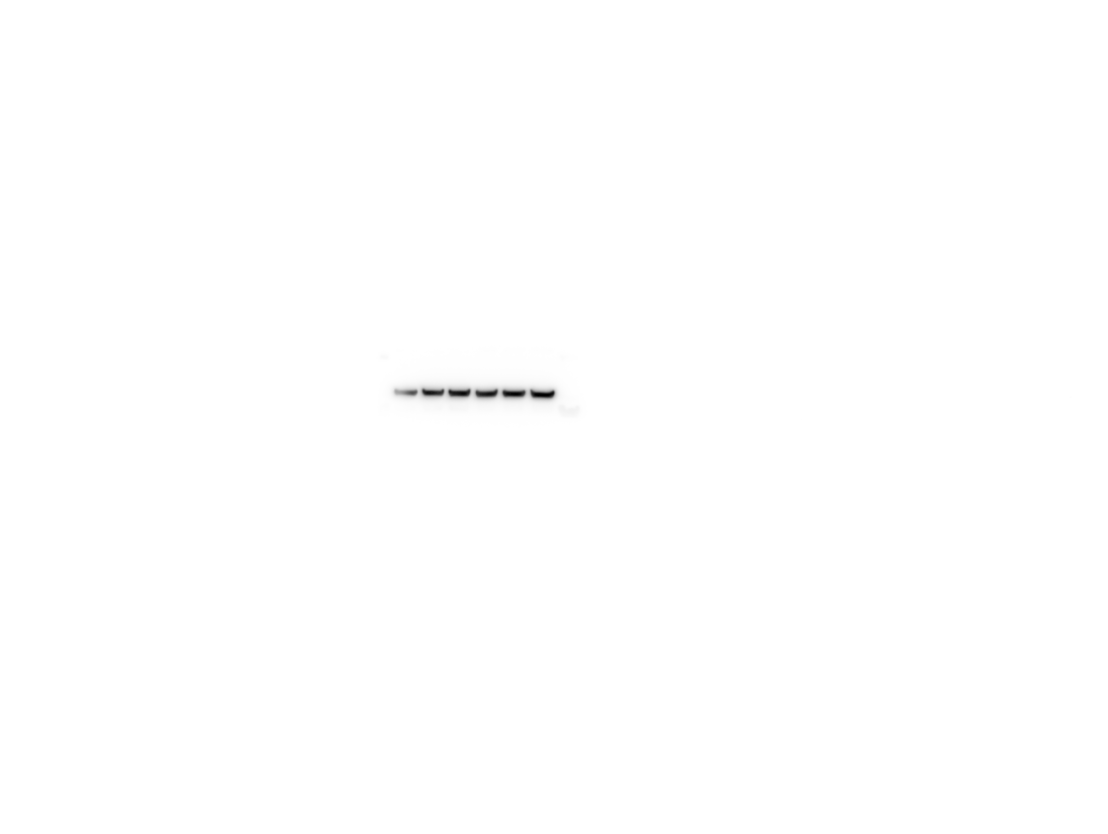

### figure_3A_source_data_hsp90_annotated.tif

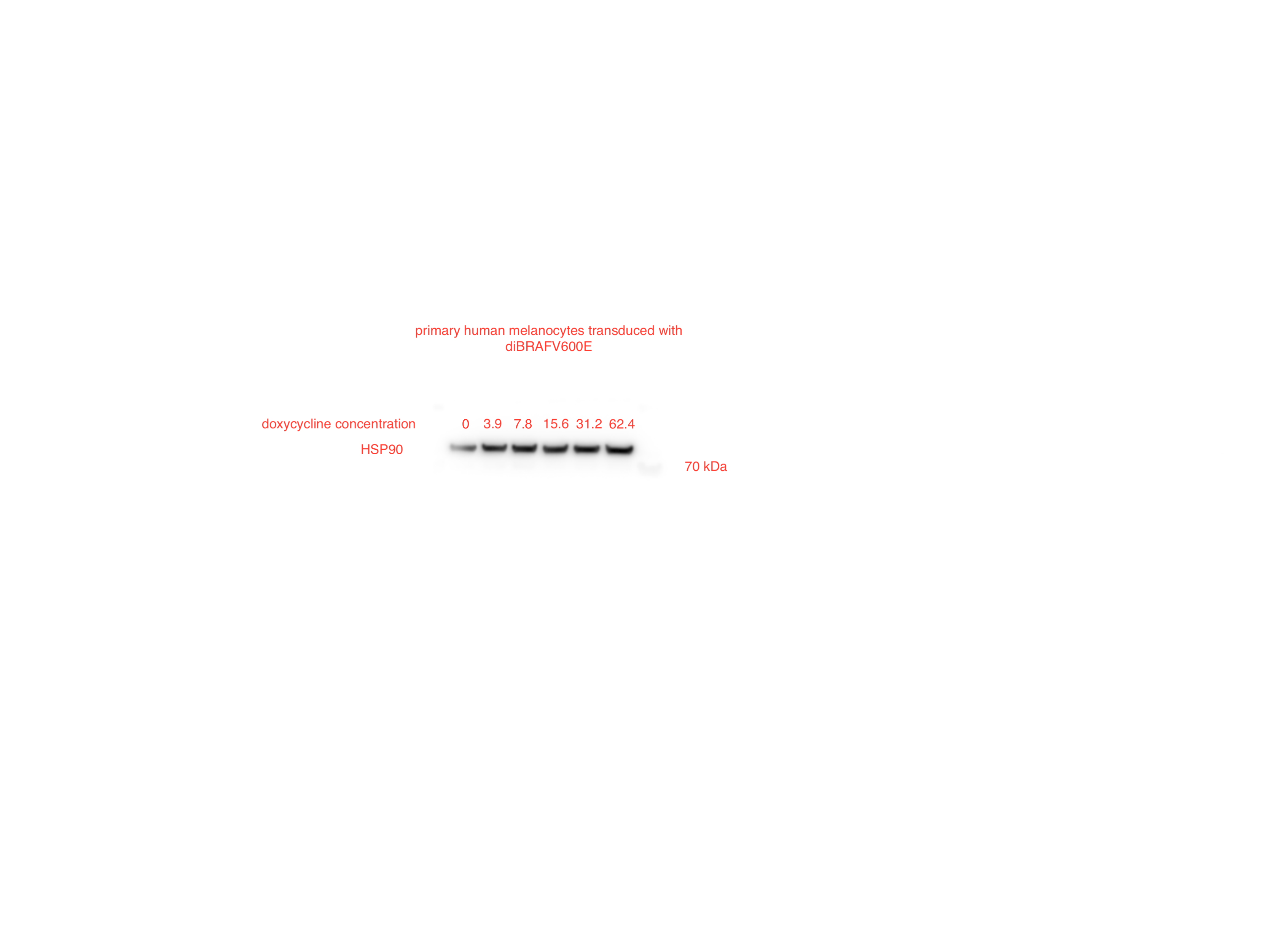

### figure_3A_source_data_phospho_erk_annotated.tiff

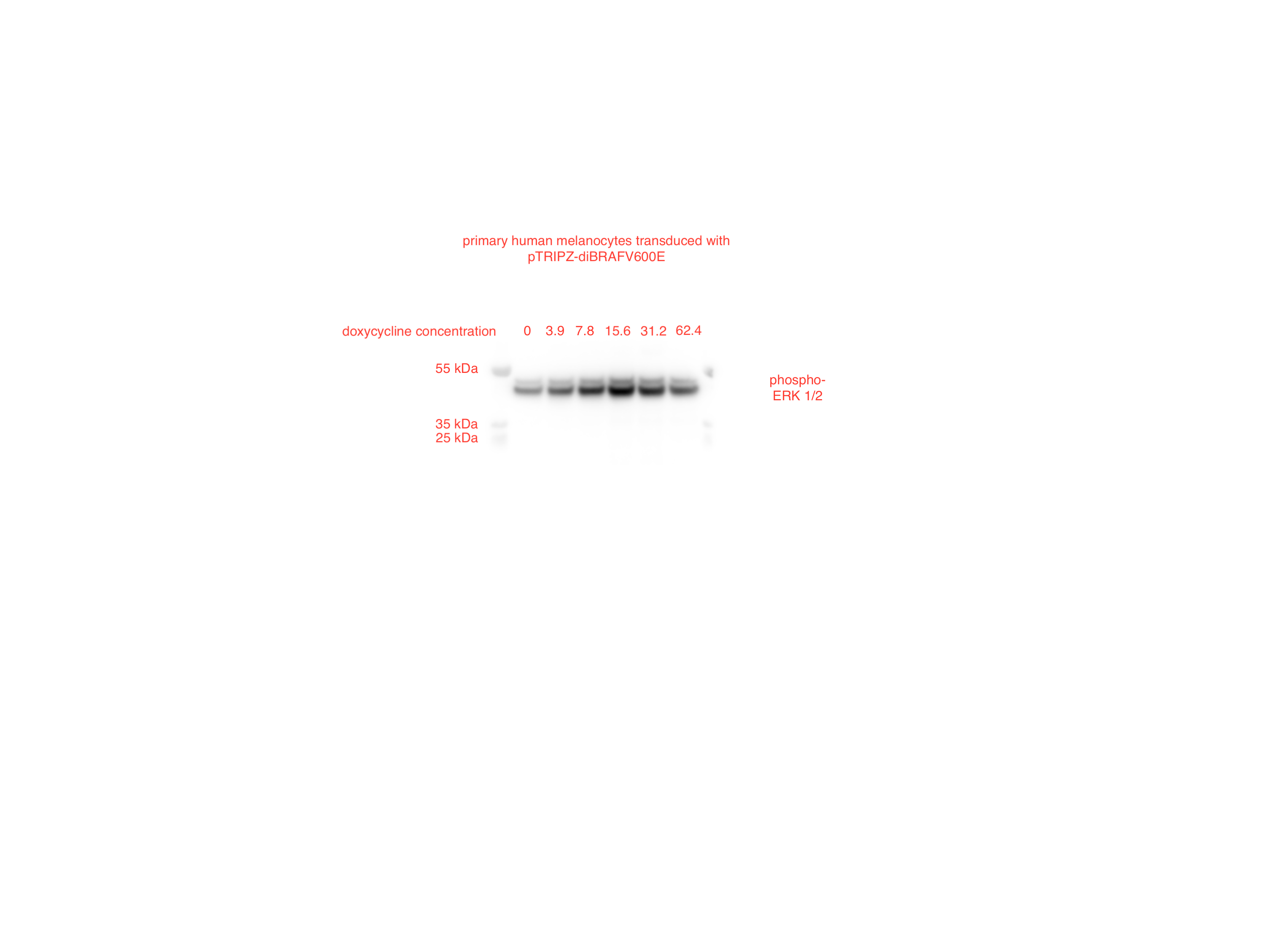

### figure_3A_source_data_phospho_erk_pub.tif

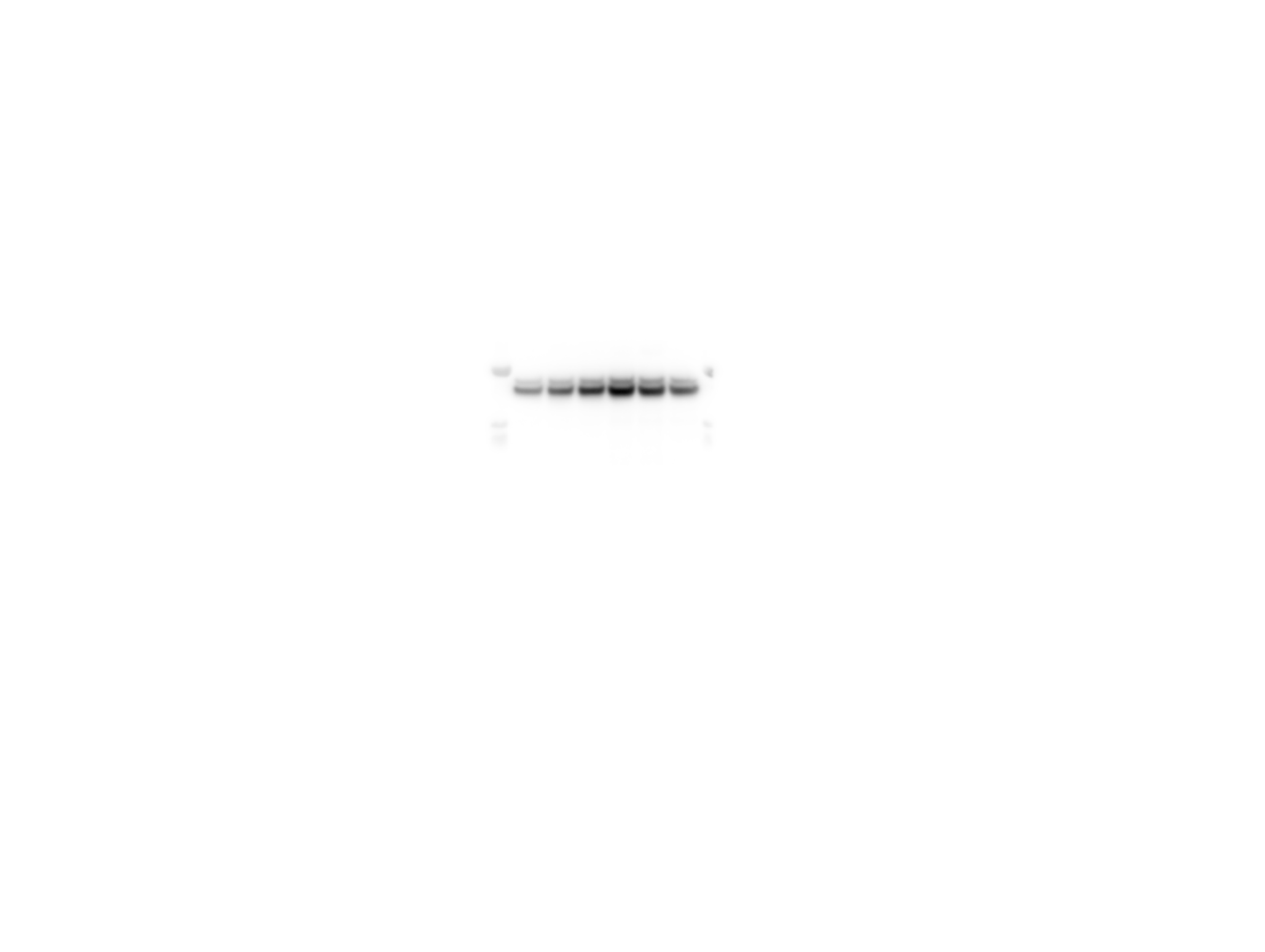

### supp_fig_S2_source_data_aurkb.tif

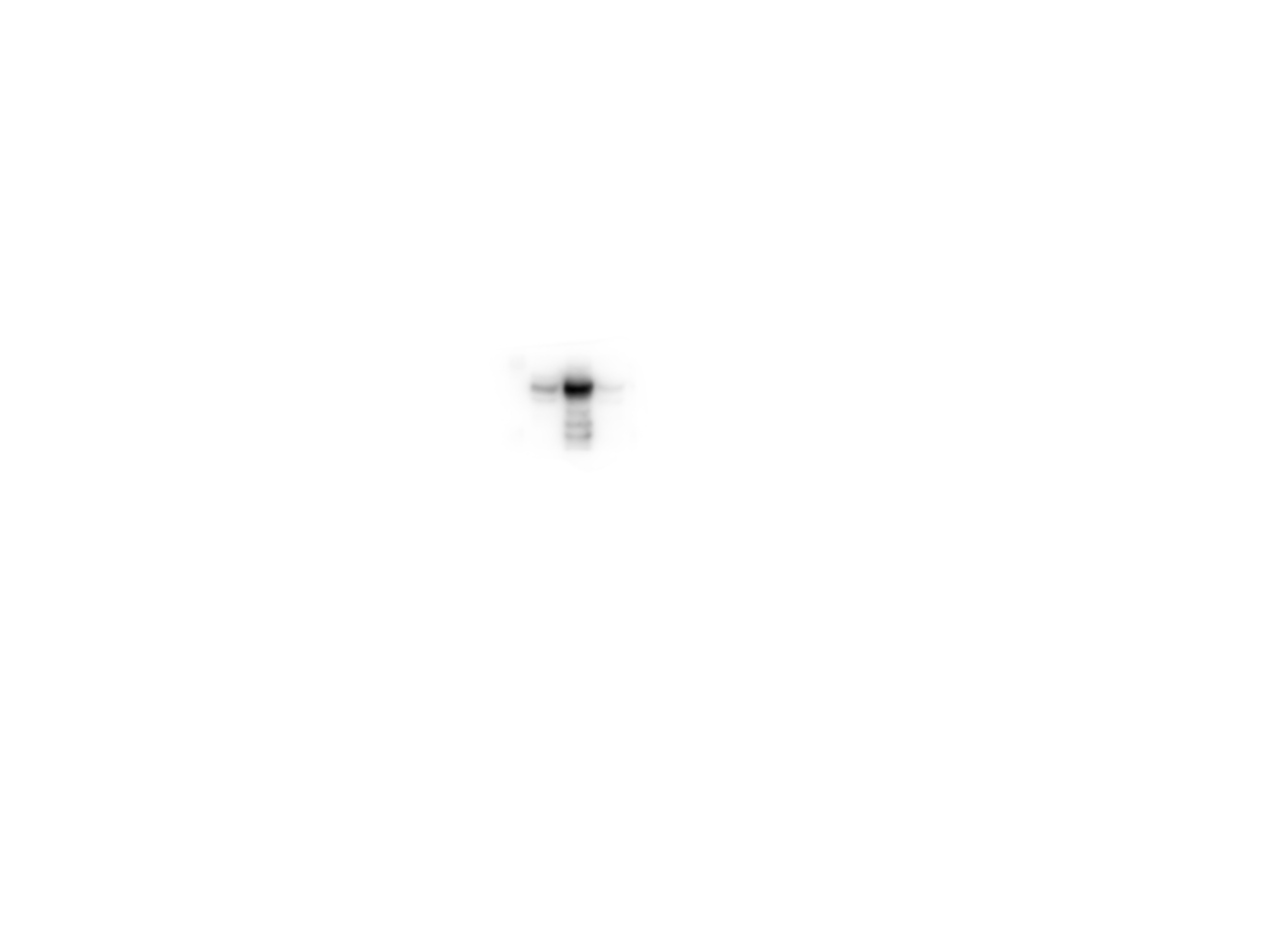

### supp_fig_S2_source_data_aurkb_annotated.tiff

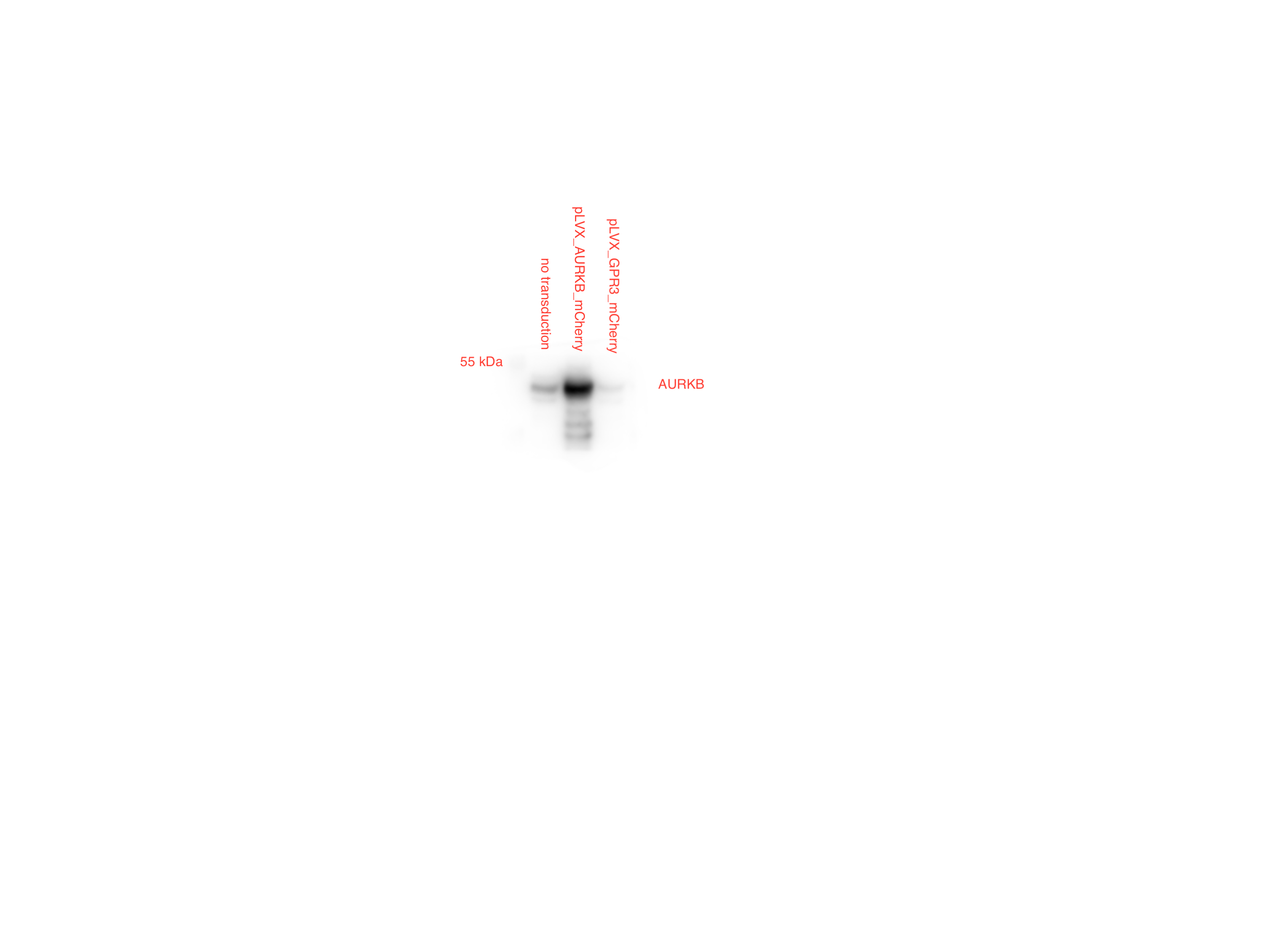

### supp_fig_S2_source_data_hsp9_annotated.tiff

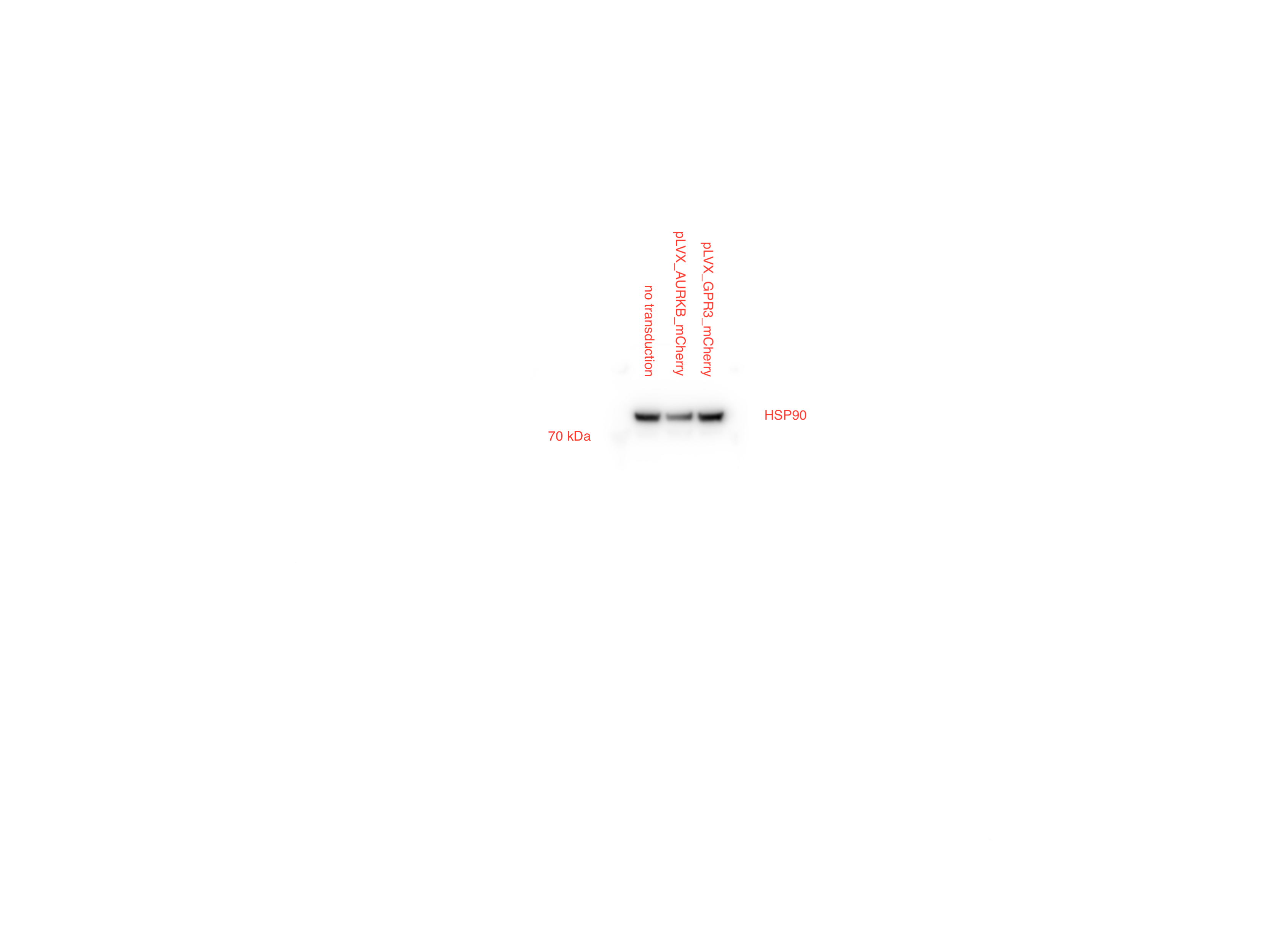

### supp_fig_S2_source_data_hsp90.tif

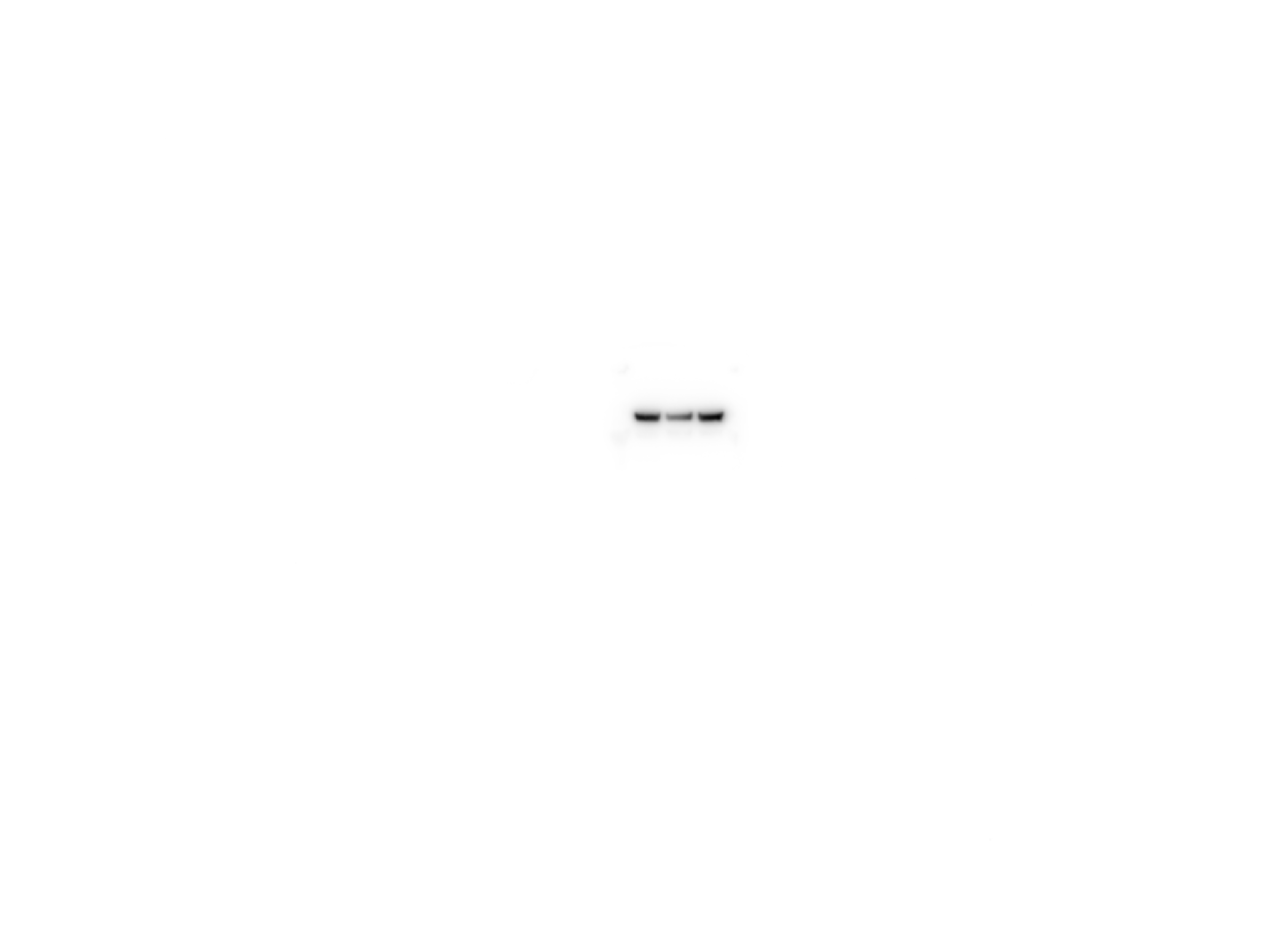
